# Estimating latent positions in social and biological networks using Graph Neural Networks in R with GCN4R

**DOI:** 10.1101/2020.11.02.364935

**Authors:** Joshua Levy, Carly Bobak, Brock Christensen, Louis Vaickus, James O’Malley

## Abstract

Network analysis methods are useful to better understand and contextualize relationships between entities. While statistical and machine learning prediction models generally assume independence between actors, network-based statistical methods for social network data allow for dyadic dependence between actors. While numerous methods have been developed for the R statistical software to analyze such data, deep learning methods have not been implemented in this language. Here, we introduce GCN4R, an R library for fitting graph neural networks on independent networks to aggregate actor covariate information to yield meaningful embeddings for a variety of network-based tasks (e.g. community detection, peer effects models, social influence). We provide an extensive overview of insights and methods utilized by the deep learning community on learning on social and biological networks, followed by a tutorial that demonstrates some of the capabilities of the GCN4R framework to make these methods more accessible to the R research community.

## Introduction

Graphs are mathematical constructs that relate entities or actors to each other and as such are well equipped to solve important issues across social network^1^, biological^2–5^, and epistemological sciences. Current research in network sciences includes developing methods to improve characterization of network structure, interpretation of how network structure relates to the behavior and characteristics of actors within the network, and how networks as entire units can be compared with one another to assess the effectiveness of interventions.

Many statistical modeling approaches, such as centrality measures, Exponential Random Graph Models (ERGMs)^6–9^ and Louvain modularity^10^, have been developed to better understand relational data. These approaches require the user to state exactly what they aim to model through selection and estimation of parameters of interest after considering a family of models with parameter spaces of varying dimensionality. This is useful for testing hypotheses about what configurations of actors and actor attributes are more prevalent in the network than expected under a simpler explanation, and through that, informing the process under which the network was generated. These tasks include, for instance, which actor a new actor will form relationships with, relationship statuses over an unobserved part of the network, and actors most likely to exit the network. In addition, specialized modeling approaches have been developed for tasks such as the separation of actors into meaningful groups (e.g. Hoff’s latent space model^11^).

However, when predicting cluster membership or attributes of network nodes, it is valuable to describe the exact means by which information is being utilized to inform the prediction. For instance, one could use a derived graph with edges weighted by the information one node passed to its partner, or sub-networks centered on each actor (ego-centric subgraph). Either approach would enumerate the precise set of higher order relationships that played a crucial role in the prediction of that neighbors’ attributes or cluster membership. From a molecular standpoint, these could be groups of proteins which interact to bring about a drug’s side effects ^12^; from the sociological standpoint, these could be groups of friends and their friends which may ultimately influence an individual to cease smoking or other addictive behaviors. The traditional statistical modeling approaches represent elegant devices for determining whether a specific phenomenon may be present in the network. However, statistical models may be less well equipped for a variety of prediction problems involving networks. As such, while traditional statistical models may provide some of these capabilities (e.g., social influence, ego-centric subgraphs), their parsimony may come at the expense of model accuracy (i.e., “All models are wrong but some are useful”), which may in turn limit the fidelity of the observations made on the network.

As such, newly developed network embedding approaches aim to assign actors in a network to a low dimensional latent position ^13–15^, where the distance between actors conveys functional relationships between actors. In contrast to previous paradigms, which emphasize network understanding over network prediction, these methods find patterns with minimal supervision and are not limited by the specification of the statistical model or heuristics. This allows the methods to more flexibly consider which information in a network is important when solving similar prediction tasks.

Deep learning represents a set of computational heuristics that searches over a large space of nonlinear transformations and interactions to arrive at an ideal model specification through the use of Artificial Neural Networks (ANN), which are inspired by nervous system processes^16^. Recent deep learning software (Graph Neural Networks; GNN) has been developed for learning patterns on graph structured data by convolving attributed information across local neighborhoods of nodes. GNN have the property of exchanging information between nodes and their neighbors to update their embeddings; for instance, the new position assigned may reflect some form of homophily (similar attributes), or heterophily (different attributes) ^17^, between a pair of actors. Subsequent applications of convolutions allow for higher order interactions between actors; for instance, there is a certain analogy here to Hoff’s latent space models and the concept of transitivity under those models. GNN have emerged as powerful means from which to analyze relational data, improving the ability to represent information between actors and across entire systems through embeddings versus traditional statistical approaches ^13,18–22^. Consequently, improvements in accuracy often come at the expense of model interpretability; however, recent model explanation techniques have sought to rectify these issues to make GNN models both highly accurate and interpretable.

While traditional statistical methods seek to identify the influential factors relating to the formation of ties by searching over explicit local network configurations and covariates, their deep learning counterparts are able to scale to larger networks and are more focused on inferring undiscovered ties or unseen networks that arise from network embeddings via an “autoencoder” design. Whereas traditional methods aim to describe the factors contributing to network structure, deep learning approaches are more focused on derivation of latent vectors and prediction of unseen dyadic relationships amongst observed actors ^21,23^.

For instance, in this paper we will consider a network data set of corporate law partnerships in the Northeastern United States, with ties indicative of perceived friendship between two lawyers^24,25^.

The following are a selection of questions one might ask of the lawyer friendship network:

- Statistical properties of the network: for instance, eigen-centrality measures, transitivity/clustering, network density, influence of actor related to seniority/age
- Key Drivers of Formation of Ties: Is friendship mutual? Is there clustering in the network? How important is being part of the same workplace? Are similarly aged lawyers friends? Exponential Random Graph Models estimate the formation of network ties using a set of sufficient statistics which define the likelihood of the graph as a whole as a function of their associated unknown parameters, whose support (parameter space) can then be searched to find the values that maximize the probability of observing the given graph conditional on the specified set of network statistics.
- Peer Effects: Do lawyers practice cases in a more similar way when they have tight network connections between them?
- Social Selection: Does seniority, schooling and law firm of lawyer influence the lawyer’s choice of practice? Does the seniority/status of other lawyers influence one’s own status (partner/associate)?
- Network Comparison: What network statistics relate to the productivity of particular law firms? What statistics provide means of comparing networks?
- Community Structure and Higher Order Transitive Relationships: Louvain modularity and latent space models are able to detect communities of actors that may be related by higher order connectivity/relationships beyond dyadic independence. If the network is dyadic independent, then there are no communities.

There are many traditional statistical and deep learning modeling approaches that may be employed to tackle these questions, but sparse literature comparing these paradigms (largely reflective of one methodology being devoted to network understanding, while the other devoted to network prediction). Table 1 enumerates types of social network analyses and examples of software from each paradigm that attempt to model each approach.

**Table 1:**
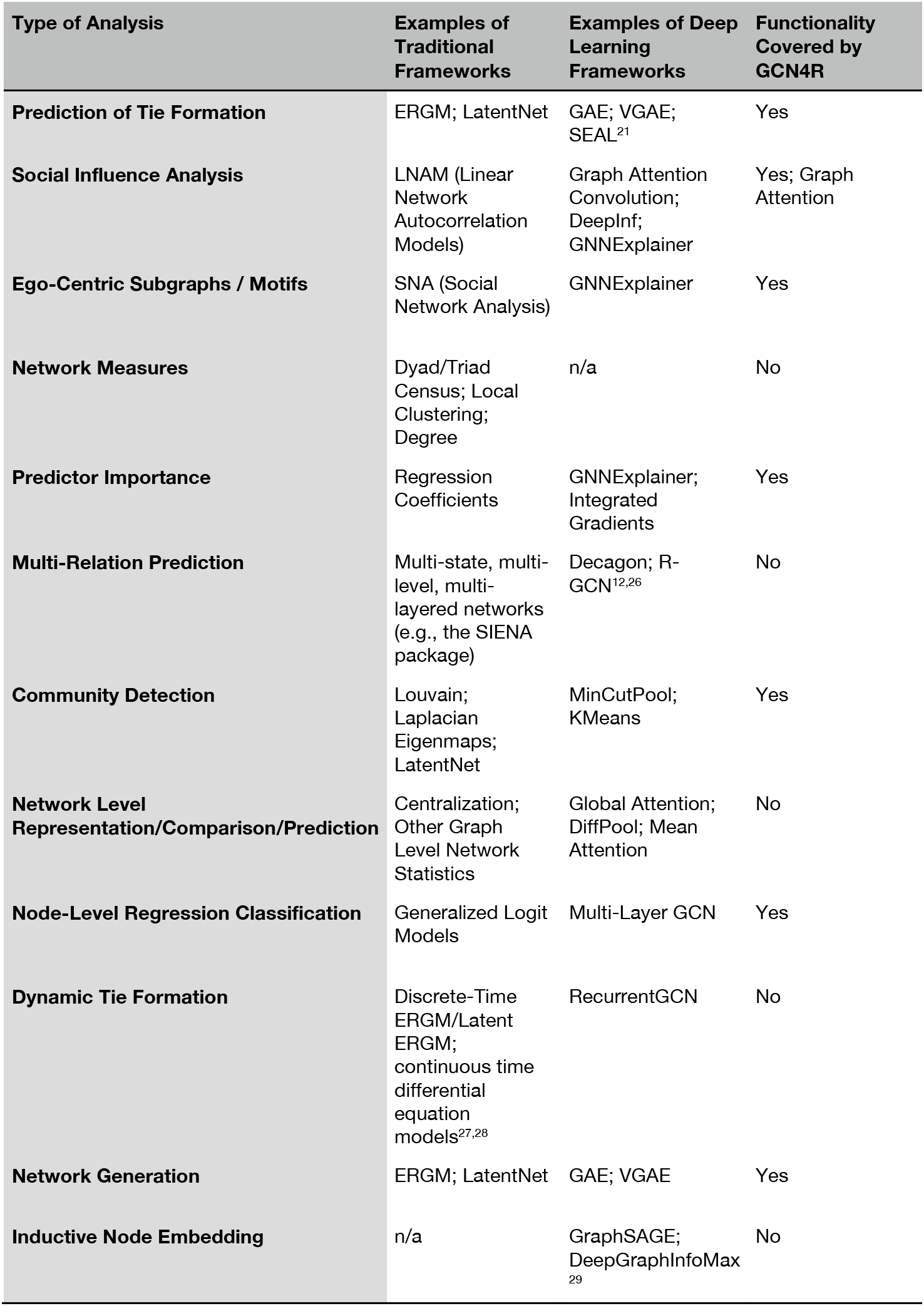
Examples of Types of Social Network Analyses and Accompanying Software Frameworks

### Contribution

Most of the aforementioned GNN software listed in Table 1 has largely remained inaccessible to researchers who only utilize R or lack a deep learning engineering background. The GCN4R package, was designed to make the class of graph neural network algorithms easily accessible to the R community, allowing for simplified model fitting procedures with the following goals in mind:

1. Easily train neural network models that are able to make predictions on graphs for the following tasks:
  a. Edge prediction
  b. Simulate new graphs
  c. Detect communities/clusters while integrating attributed information
  d. Predict node-level characteristics (classification and regression; imputation of missing characteristics of observed nodes)
  e. Form node embeddings / network layouts to supplement other network software
2. Summarize the prediction results and model diagnostics; visualizations that inform the results from the estimating procedures
3. Interrogate important nodes, edges, attributes and specific structural motifs as determined by the neural network

The GCN4R package contains functions and classes that wrap and extract results from neural network fitting procedures in Python, under the PyTorch Geometric deep learning framework ^20^. These functions make fitting and interrogating graph neural networks easy to execute after the data has been loaded, with complete functionality in R. In Table 1, specifies of the social network analysis methods included in GCN4R are presented.

The GCN4R package is available through GitHub via the repository jlevy44/GCN4R, which can be installed using devtools. GCN4R has 23 package dependencies, and help documentation is available for each of the R functions or through a Wiki page and demo available on our GitHub repository and linked to this paper (https://github.com/jlevy44/GCN4R/wiki). Below we discuss the capabilities of GCN4R and some of the underlying theory and key contributions, and demonstrate its functionality through an illustrative example.

## Theory

### Graphs

A graph consists of entities (vertices/nodes, *V*) and the enumerated relationships between the entities (edges, *E*) ^30^. Of particular interest is the study of dyadic relationships, which are captured through a list of edges. Directed graphs explicitly state the directionality of these dyadic relationships, where one node is the sender, and the other the receiver. In undirected graphs, a particular edge may be unordered. The set of vertices and edges constitute a graph, *G*=(*V,E*). This mathematical structure can also be constructed using an adjacency matrix, ***A***, a square matrix of which the vertices constitute the rows and columns, while an edge (*i, j*), corresponding to nodes *i* and *j*, is an indicator function that is valued 1 when there exists a relationship between *i* and *j* and 0 otherwise. The adjacency matrix is endowed with a number of properties that can be conveniently expressed using linear algebra, which in turn streamlines computations on graphs. When the adjacency matrix is a sparse matrix, calculations can be streamlined by expressing them (or approximations thereof) directly in terms of the edge list. Nodes may be attributed by a design matrix, 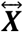, with node-level covariates.

Throughout the rest of this paper, we will define 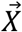 as the data that is input into the model, *W* a matrix that describes learnable model parameters/weights/kernels, 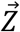 a node-level embedding after applying transformations to 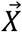 and ***A***, and 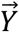 the model’s classification or regression prediction.

### Neural Networks

ANN represents the predictors of the aforementioned covariate/design matrix as a set of nodes, or an n-dimensional vector 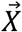 for a single observation. This information is passed to multiple hidden layers of nodes that combine and transform the information from the previous layers of nodes into a compressed representation 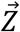. The ANN as a whole maps 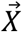 into a low dimensional representation 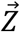 via the functional *f* : *x* → *z*, typically of the form *f*(*x*) = *σ*(*Wx* + *b*) which can represent some target outcome or representation of interest (σ may be a nonlinear function such as the logistic, hyperbolic tangent, rectified linear unit, or identity functions). Multi-layer perceptrons (MLP) are used to transform data that are represented by vectors (1-rank tensor; where tensors are generalizations of scalars, vectors, and matrices to a possible greater number of components), while convolutional neural networks (CNN) slide learnable filters across images (matrix or multidimensional array; 2-3 rank tensor) to extract and integrate lower order structural and color motifs into higher order abstractions to make a prediction ^31^. Various objective functions (*L*) such as mean squared error 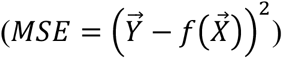 are utilized to update the parameters of these models via backpropagation.

### Overview of Graph Neural Networks

While MLPs and CNNs may be applied readily to various biomedical data, such as images and gene expression profiles, these methods are not easily extended to network data. While convolutions over images propagate information within a fixed neighborhood of pixels and require consistent ordering of predictors, graph neural networks relax the convolutional operator used for image analyses (a parameterized grid structured kernel) to aggregate information across an unfixed number of neighbors as denoted by the adjacency matrix ^20^.

In the image analysis domain, parameterized convolutional kernels *g* (actualized by kernel matrix *W*) slide across 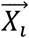, where 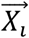, is an image, to yield a “feature map” *Z* via the operation:

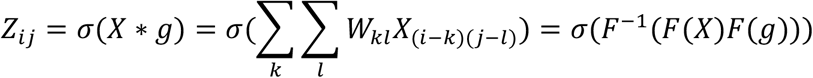

The convolution is equivalent to multiplying two signals together in their frequency space via the Fourier transformation *F*, then inverse transforming via *F*^−1^ to yield the convolved signals. In its simplest formulation the graph convolution may be expressed similarly, where the latent embeddings of the nodes are unnormalized:

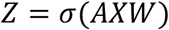

Here ***A*** is a fixed *n* by *n* adjacency matrix, *X* is a *n* by *m*_1_ covariate matrix, *W* is a parameterized *m*_1_ by *m*_2_ weighted projection matrix, and *Z* is an *n* by *m*_2_ embedding matrix. This equation has the property of summing up the predictor vectors of the neighborhood nodes and transforming their dimensionality from *m*_1_ to *m*_2_ to generate latent embeddings, *Z*, for all *n* individuals. Many alternative specifications of signal propagation across a network have spawned from this early formulation.

The more popular expression of the graph convolution is the “message-passing” formulation, which directly relates how information is propagated in a network through successive aggregation of node-level predictors across edges of the graph. Here, we discuss how information for one single node is updated for one convolutional layer:

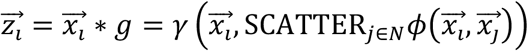

The predictors of the neighbors, *N*, of node *i* are represented by 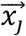. A differentiable function/neural network, *ϕ*, which may contain learnable parameters, maps 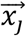 to 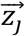. The operations performed on the neighbors of node *i* to update their information is done by scattering option SCATTER, which parallelize executions across the CPU/GPU (sending “messages” outwards and gathering the results). Since the ordering of the nodes does not matter for updating the embedding of 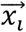, a differentiable function *γ* aggregates the hidden state information of the neighbors of *i* (and if there are self-loops, of node *i* itself) via a sum or weighted average-like operation. For instance, *ϕ* could be a bilinear projection of 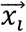 and its neighbor 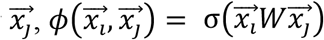, while *γ* could be the averaging operation, 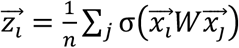 for *n* neighbors. In this way, information from the neighbors *N* of 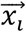 are propagated to node *i* to update its embedding to 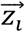. Multiple applications of these convolutional layers expand the neighborhood from which information is propagated over (since the updated embeddings of the neighbors contains information passed from their neighbors), to consider higher-order dependence between the individuals in the network.

These filters contained within each graph convolution may generalize to unobserved/unseen graphs to update node-level latent embeddings in a similar manner, though these capabilities are not yet offered in this framework. Graph-level summaries (one vector per graph) may be formed through aggregation/pooling operations (AGG; e.g., mean, agglomerative clustering centroids) across the nodes of the network to compare multiple graphs to one another:

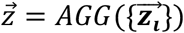

### GNN Model Formulation with GCN4R

The GCN4R package operates on single graphs in the R environment. The user selects a backbone convolutional operator, which sets *γ* and *ϕ*. In this case, the operator is the GCNConv^32^. Multiple applications of the operator update the node-level predictors to form latent embedding 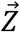 for the nodes in the graph. Like the latent space models utilized by software packages such as ‘*latentnet’*^8^, the distance between actors in these embeddings represent higher-order dependencies. Similarly, the distance between the actors may be indicative of the presence of an edge. Therefore, the outer product of the embeddings may be used to approximate the original network. These methods constitute what are known as the “auto-encoder” methods, where an encoder maps the node-level covariates of the input graph to low-dimensional representations, then decodes this information to attempt to recapitulate the original graph structure, and outputs a probability of an edge as such: 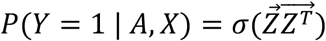. Alternative decoders may be specified which correspond to latent space and latent distance models, given by 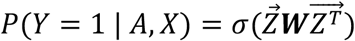 and 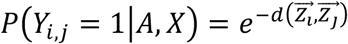 respectively, where *d* is the distance between two latent actors and σ is a sigmoid or logistic function^33,34^. If we consider the formation of each edge as a Bernoulli trial, the primary objective of such a model is to maximize the likelihood of these Bernoulli trials, which we refer to under the machine learning specification as the reconstruction loss: *L_R_*, alternatively referred to as the likelihood function by the statistical community. Note that these probabilities may vary across all edges of the graph (i.e., not an Erdős–Rényi model). The latent vectors 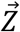 may also be assumed to follow a multivariate normal distribution, emulating the assumptions from stochastic variational inference (SVI)^35^. The distribution of the latent vectors is the posterior distribution *P*(*Z*|*X*), where the decoder represents the likelihood function or data generating process given the latent data *P*(*X*|*Z*). The Kullback Leibler Divergence, *L_KL_*, is minimized between the distribution of latent variables and the “guide” multivariate normal distribution to approximate the true posterior distribution in the absence of a conjugate-prior. Alternative specifications of the KL divergence to match a prior distribution is the adversarial regularization^23^, which uses a discriminator as a function to approximate the densities of both the prior and generated latent distribution and attempts to decipher whether the latent sample came from the prior or generated posterior sample, the loss of which is *L_adv_*. Regularization, for deep learning models, is the introduction of a penalty term into the loss function to discourage model complexity; in this case, to constrain the shape of the posterior distribution. Additional penalty terms alters the objective function for fitting the model and, therefore, the predictions of held-out node covariates (mean-squared error or multinomial likelihood maximization / cross entropy loss may be applied for node-level regression and classification respectively; *L_pred_*). Finally, there exists additional specifications for community detection, where the objective may be represented by loss functions that are related to K-Means clustering^36,37^ (minimize distance to cluster center) or Laplacian eigenmaps^38^ (min-cut pooling; minimize the number of links to prune in a network; identify information diffusion weak points), *L_cluster_*. Each loss function may be weighted by a hyperparameter *λ*, to form the final modeling objective:

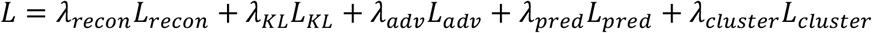

As in general, once the likelihood function has been specified, optimal estimators of the unknowns (e.g., node attributes and edge-statuses) are then defined (often implicitly) and their values on a given data set become the target of the computational algorithms used to implement the estimation method.

## Interpreting Results

Once a GNN model has been fit, the GCN4R package offers some preliminary means of interrogating edges, nodes and higher-order structural motifs that had an important impact on the predictions. Some of the provided capabilities are popular methods in the deep learning community for training and interpreting graph neural networks. Here, we outline some of the theory behind three interpretation methods ^39^: attention for edge-wise importance, integrated gradients for uncovering important covariates, GNNExplainer to extract ego-centric motifs and important node-level covariates.

### Attention to Uncover Important Edges

Attention mechanisms in deep learning are motivated by our own visual attention processes which allow us to focus on the most important regions of an image or scene for classification tasks, ignoring irrelevant or redundant information. This mechanism may be codified by a vector of importance scores for a given observation which upweights the contribution of that particular aspect of the data for any learning task. While there are various attention mechanisms that may be applied over graphs, our package utilizes the “graph attention” mechanism via selection of the “GATConv” operator^40^. The graph attention convolution operator calculates for each node an importance score for its surrounding neighborhood:

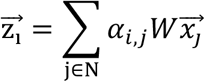

Here *W* serves as a linear operator to project 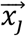 into another subspace and is updated during the training process. While the graph convolution can be seen as an averaging of the latent positions of the neighborhood of node *i*, the attention mechanism learns to weight the edges between each of the neighbors *j* and node *i*, (*α_i,j_*) to alternatively specify a weighted average of the positions. The node in the neighborhood with the highest attention weight is said to most strongly influence or contribute to the latent position of node *i*. Given multiple applications of attention layers, the attention for each layer may be found to be different and thus may relate to the degree of some higher order dependency between actors.

### Integrated Gradients for Establishing Important Predictors

Important features for each learned model can be assessed using backpropagation techniques such as integrated gradients. While these neural network approaches may yield highly accurate models, they are often thought of as “black boxes”, from which input data is manipulated by this box to yield an output without an understanding of how the box works. One traditional point of inspection is calculating the gradient of the learned model for a particular observation, 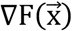, which serves as a first order approximation of the non-linear fit. The importance of each predictor to the prediction is then 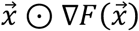 (denoting elementwise multiplication). However, the gradient saturates at extreme values and has no defined baseline for comparison, leading to inconsistent attributions of predictors. As such, perturbation methods were developed to learn about how the model arrives at its prediction though perturbation of the input features, which ultimately changes the model’s prediction. These changes can be back propagated through the network to reveal important predictors. Integrated Gradients (IG) acknowledges the presence of a baseline, then calculates and sums more informative gradients on the path between this baseline 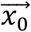 and the observation 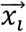 given a series of increasing perturbations, encapsulated in the below formula^41^:

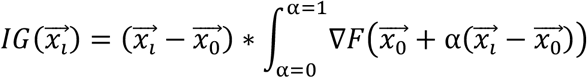

The integration of these backpropagated changes in prediction (predictor importance) across the sequence of these perturbations to yield final importance scores for predictors. Predictor importance may be found for individual observations and aggregated across the nodes.

### Egocentric Motif Extraction and Predictor Importance

While importance of dyadic dependence and various predictors may be assessed through attention and integrated gradients, there exist richer higher order structural motifs / egocentric subgraphs that contribute to a node’s prediction. For each node, GNNs pass information from the node’s *k*-step neighborhood (acquired after applying *k* convolutions to the network and similarly to cross-sectional network autocorrelation models with AR(k) dependency). In light of this, model explanations for node or graph-level predictions may be provided in the form of a learned egocentric subgraph that correctly identifies the information pathway. The GNNExplainer learns which subgraphs/motifs and features are most important for node-level predictions (e.g. for some nodes, this could be a subset of second-degree neighbors, while for others a subset of their third- or fourth-degree neighbors; or a subgraph of complete *k*-step egocentric subgraph, or some other subset). In general, there is no way of sampling a network such that all of the structure in the original can be retained, unless the network has a simple model^42^. The GNNExplainer learning task is operationalized by learning a mask over the graph (weighted adjacency matrix which may be thresholded; which serves to prune irrelevant nodes from what would otherwise be an egocentric subgraph in traditional network models), and predictors (vector of predictor importance; prunes irrelevant predictors), that maximizes the mutual information between the subgraph, its selected predictors, and makes the prediction with:

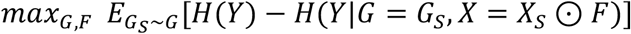

The subgraphs *G_S_* are represents a family of plausible graphs, whereas *F* represents a learnable vector of indicators valued between 0 and 1 that effectively prunes predictors. The learned mask, *M*, that exists over the adjacency matrix *A*, may take on values between 0 and 1 over true edges. A threshold may be specified over this mask to convert it into an indicator matrix to effectively prune the edges from the learned subgraph that the model feels are less important to the node’s prediction.

### Perturbation Methods for Assessing Significance of Important Predictors, Edges and Motifs

Formal methods for assessing predictor, edge, and motif importance from GNN interpretation methods have not been established. Here, we provide a general framework for filtering model results to reveal important associations. First, the original graph *G*_0_ is passed through an interpretation framework to yield importance scores *IS*_0_ (e.g., *α_i,j_* or *M*). Then, many graphs, *G* = {*G*_1_, *G*_2_, …, *G_N_*}, (on the order of one hundred) are generated using either: 1) the Erdős–Rényi model under a fixed edge probability *p*; or 2) may be perturbed from the original graph, where positive edges are randomly removed with probability *p_pos_* and negative edges are randomly flipped to positive edges with probability *p_neg_* and can be set such that the density of the network is preserved. When these graphs, *G*, are passed through the GNN, it is expected that many importance scores *IS* (e.g., *α_i,j_* or *M*) from the previous sections will decrease. The interpretation framework maps each graph *G_i_* to importance score *IS_i_*. The original importance score *IS*_0_ can then be compared to the importance scores from the generated graphs: *IS* = {*IS*_1_, *IS*_2_, …, *IS_N_*}. A one-sample Wilcoxon signed rank test can assess the following hypothesis tests for each predictor, edge or motif to assess their significance:

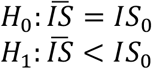

The calculated p-value can be adjusted using a post-hoc adjustment for multiple comparisons to reduce Type I error (e.g., Bonferroni). If generating graphs under the Erdős–Rényi model, statistical significance is with respect to the comparison between the current graph and generated graphs under the Erdős–Rényi model (i.e., is a test of whether the graph is the random graph against the alternative that some other model generated the graph), while for perturbed graphs, significance indicates how sensitive/stable the importance score is to perturbation of key information pathways (i.e., the test is tests whether the observed graph has features beyond those that are preserved by the perturbation). In the example given earlier, if *p_pos_* = *p_neg_* then the density of the network is being preserved but any other structure in the network is removed by the randomization.

### Variational Methods and Alternative Model Optimization Schemes

The GNN modeling approach introduced in GCN4R only covers a small subset of all possible network learning tasks using GNNs. Here, we outline a few other potential possibilities that may be accomplished with our framework.

This framework may also be configured for the generation of new graphs. The latent parameters as introduced by the encoder-decoder architecture of the GNN follows a variational bayes learning framework^43^, where graphs are generated via some data generating process. The likelihood function is a multivariate Bernoulli distribution given by the decoder:

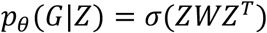

The prior, *p_θ_*(*Z*), of the data generating process is assumed to be a multivariate normal distribution. The evidence/marginal likelihood is intractable to compute and is estimated using the evidence lower bound (ELBO). Variational inference may thus be utilized to approximate the true posterior since it cannot actually be derived. The model/encoder *q_ϕ_*(*Z*|*X*) approximates the posterior distribution *p*(*Z*|*G, X*) through matching to a known family of distributions by minimizing the KL-divergence:

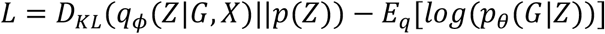

This penalization serves to emulate conjugate Bayesian analysis by finding the closest approximating distributions such that a conjugate relationship holds. These variational principles are related to the Calculus of Variations, which is a technique that identifies the best function after searching over a space of plausible functions (i.e., changing a function and deciding whether fit has been improved). In this case, the search is for the “best” approximate posterior distribution in the presence of an intractable integral.

As aforementioned, we weight the KL-divergence using a *λ_KL_* tuning parameter and incorporate the right-hand term as the reconstruction loss, defined to be the negative log likelihood of a multivariate Bernoulli distribution. By weighting the KL divergence higher, we can enforce the stronger constraint of having the latent vectors follow a multivariate normal posterior. We increase the generative properties of the decoder by sampling this lower dimensional distribution.

Other alternative modeling approaches that may be utilized in the GCN4R package include node-level classification and regression, denoted mathematically as:

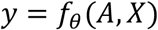

Here the model may be trained using a loss function that is the negative log likelihood of a multinomial outcome for classification tasks, or the mean squared error (MSE) for regression tasks. When training and predicting on a single graph, or setting aside nodes for testing from other graphs, we refer to these tasks as “semi-supervised” tasks^32^, where outcomes from actors within a network are propagated to their higher-order neighborhood.

## Implementation in R

### Package Installation

The GCN4R package may be installed and loaded via the following commands.

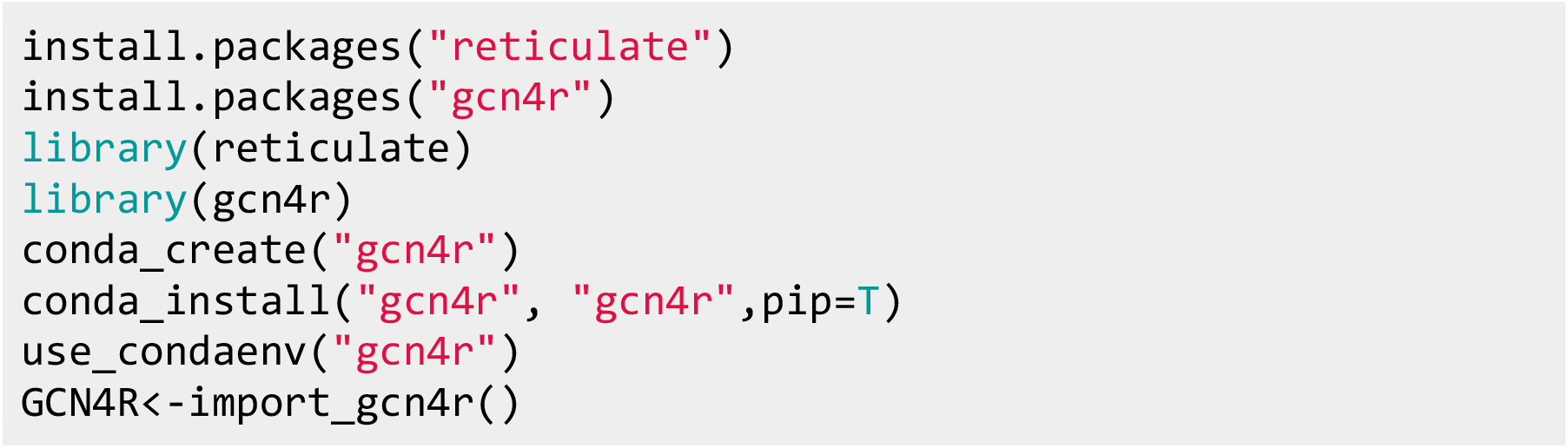

The GCN4R package requires the use of reticulate, which internally wraps and executes python subroutines built into a related python package, also titled GCN4R. The python package accesses graph neural network subcomponents using the PyTorch Geometric framework, based on the deep learning library PyTorch.

## Illustrative Example

As a use case for our package, we investigate a network of corporate law partnerships in the Northeastern United States^24,25^. The aim of the original study was to study cooperative relationships amongst 71 lawyers that had formed amongst competitive law firms. Three different networks were featured in this study: one defined by whether advice was received by individuals in a network (advice network; directed), the second by whether the lawyer considered another lawyer a friend (friendship network; directed), the final by whether two individuals were direct coworkers (coworker network; undirected). Each node was assigned attributes based on their age, gender, practice (whether they were a corporate or litigation lawyer), status (partner or associate of firm), seniority in company (number of years) and law school. Throughout the remainder of this paper we analyze the friendship network.

Here, network data from the Lazega friendship network was loaded and categorical variables were transformed into factors that build a design matrix (Table 2):

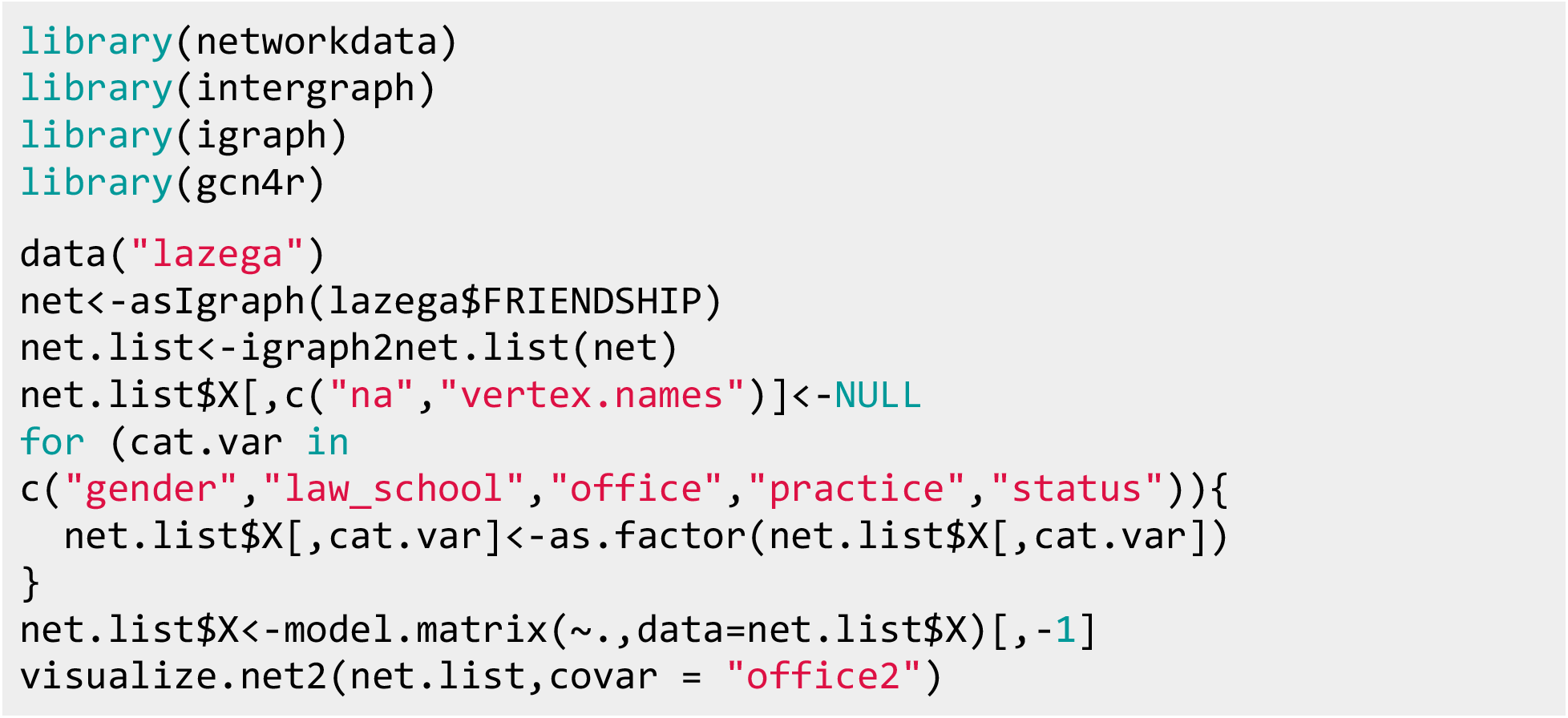

**Table 2:**
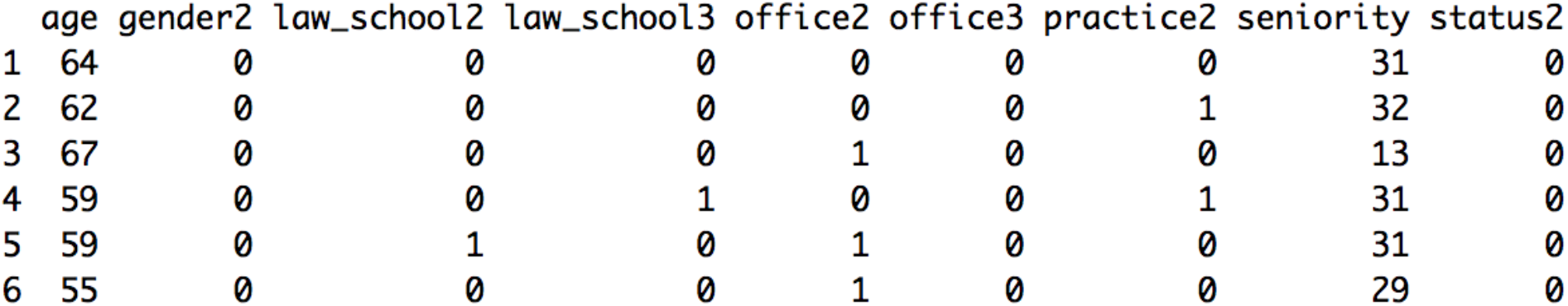
Visualization of first six actors of design matrix; gender2 indicates whether actor is male; law_school2 indicates prior attendance of the second law school; office 2 indicates actor works in second law firm; practice 2 is an indicator if the actor works in litigation; status 2 indicates whether actor is an associate of the firm

Upon initial inspection, the formation of a friendship appears to be heavily influenced by whether a lawyer worked at the same practice or was a part of the same firm.

While many of questions regarding the Lazega network can be addressed by the suite of statistical modeling tools including those outlined in the above section, the approaches present user-provided model specifications of the predictors to the outcome of interest. Through this tutorial, we aim to highlight data-driven insights of properties of the network lying outside of those obtained using traditional modeling approaches. We will primarily focus on community detection within the Lazega network using graph neural networks as an explicit use case and highlight additional functionality that estimates peer effects (node level classification and regression) and the ability to simulate/generate networks. A comparison between methods accessible between the deep learning and statistical frameworks may be found in the section: “Additional Comparisons to Traditional Approaches”.

### Fitting a Node Partitioning Model

GCN4R parameters includes many tunable default parameters, including information about how long to burn in the clustering and KL losses, neural network architecture (number of nodes, layers, backbone), number of training iterations, clusters to form, learning rate, amongst others. Essential GCN4R parameters and default specifications are shown in Table 3.

**Table 3:** Reference, definitions, and defaults of important parameters for model fitting procedures

| Hyperparameter | R Variable Name | Description | Default Value |
| --- | --- | --- | --- |
| **Learning Rate** | learning_rate | Overall step size of parameter update after calculation of gradients | 1e-4 |
| **Number of Epochs** | n_epochs | Number of times to update GCN model parameters | 300L |
| **Convolutional Operator** | encoder_base | Decides which convolutional operator to apply (e.g., Graph Attention instead of unweighted updates) | 'GCNConv' |
| **Number of Nodes Per Hidden Layer** | n_hidden | The covariates will be transformed to this dimensionality after application of each convolution | 30L |
| **Number of Hidden Layers** | n_layers | Increasing the depth of the neural network | 2L |
| **Architecture of Discriminator** | discriminator_layers | Specifies the shape (length of list is number of layers; each element is number of nodes) of the multi-layer perceptron to decide whether posterior sample is similar/different from prior/guide | c(20L,20L) |
| **Type of Autoencoder** | ae_type | Choose one of four encoder-decoder architectures; there exist variational and adversarial variants that activate their respective loss functions | 'GAE' |
| **Whether to Include Bias Term** | bias | Whether to add a learnable offset at each layer | T |
| **Weighting of KL Divergence Loss** | lambda_kl | How much to match latent vectors to multivariate normal? | 1e-3 |
| **Weighting of Adversarial Loss** | lambda_adv | See above | 1e-3 |
| **Weighting of Cluster Loss** | lambda_cluster | Emphasis on clustering nodes in latent space via K-Means or Spectral Clustering | 1e-3 |
| **Weighting of Reconstruction Loss** | lambda_recon | Emphasis on recapitulating original graph via decoder | 1. |
| **Weighting of Prediction Loss** | lambda_pred | Emphasis on estimation of peer effects (node classification/regression) | 0. |
| **Burn-In Time Before Start Clustering** | epoch_cluster | Number of training iterations before introducing the clustering loss | 301L |
| **Burn-In Iterations Before Start of KL Loss** | kl_warmup | Number of iterations before introduction of the KL divergence loss for training stability | 20L |
| **Number of Clusters** | K | Number of clusters to fit via KMeans or Spectral Clustering | 10L |
| **K-Means Clustering Iterations** | Niter | Number of iterations to optimize KMeans clustering | 10L |
| **Use Spectral Clustering instead of KMeans** | use_mincut | Whether to utilize Spectral Clustering to inform cluster assignment and regularization on model | F |
| **Column of Design Matrix to Predict** | prediction_column | Selection of a column of design matrix to perform classification/regression on | -1L |

These parameters may be generated using the following command:

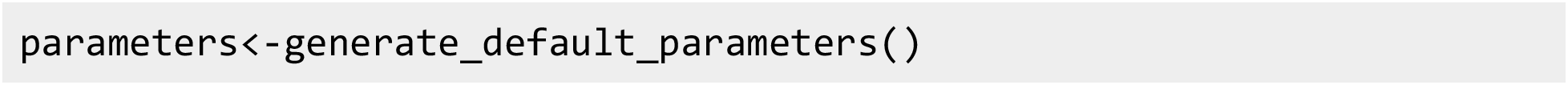

Alternatively, these parameters may be updated as:

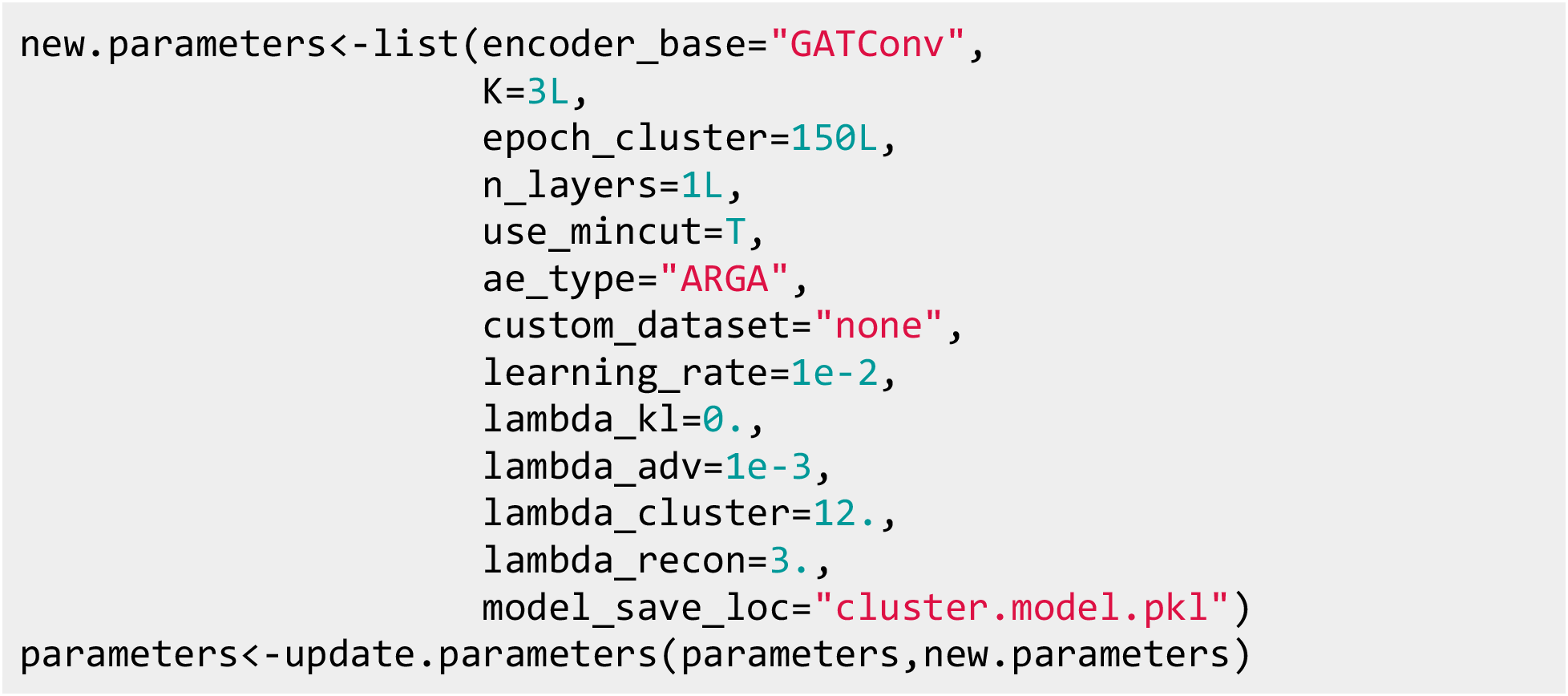

Finally, once the parameters for the learning approach have been chosen, the model is ready to fit. Here, we aim to detect up to three communities in the Lazega Network:

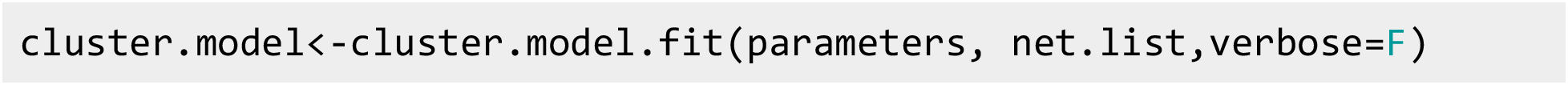

## Fit Summary and Diagnostics

Clustering results, as well as the model diagnostics and summary may be printed after fitting the model (Figure 3A):

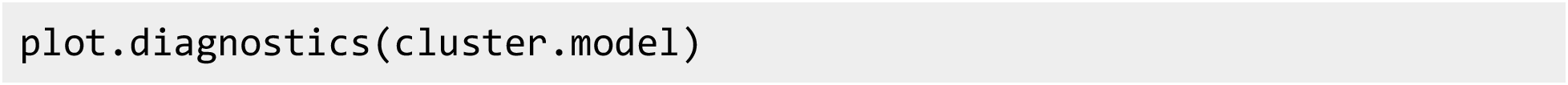

**Figure 1:**
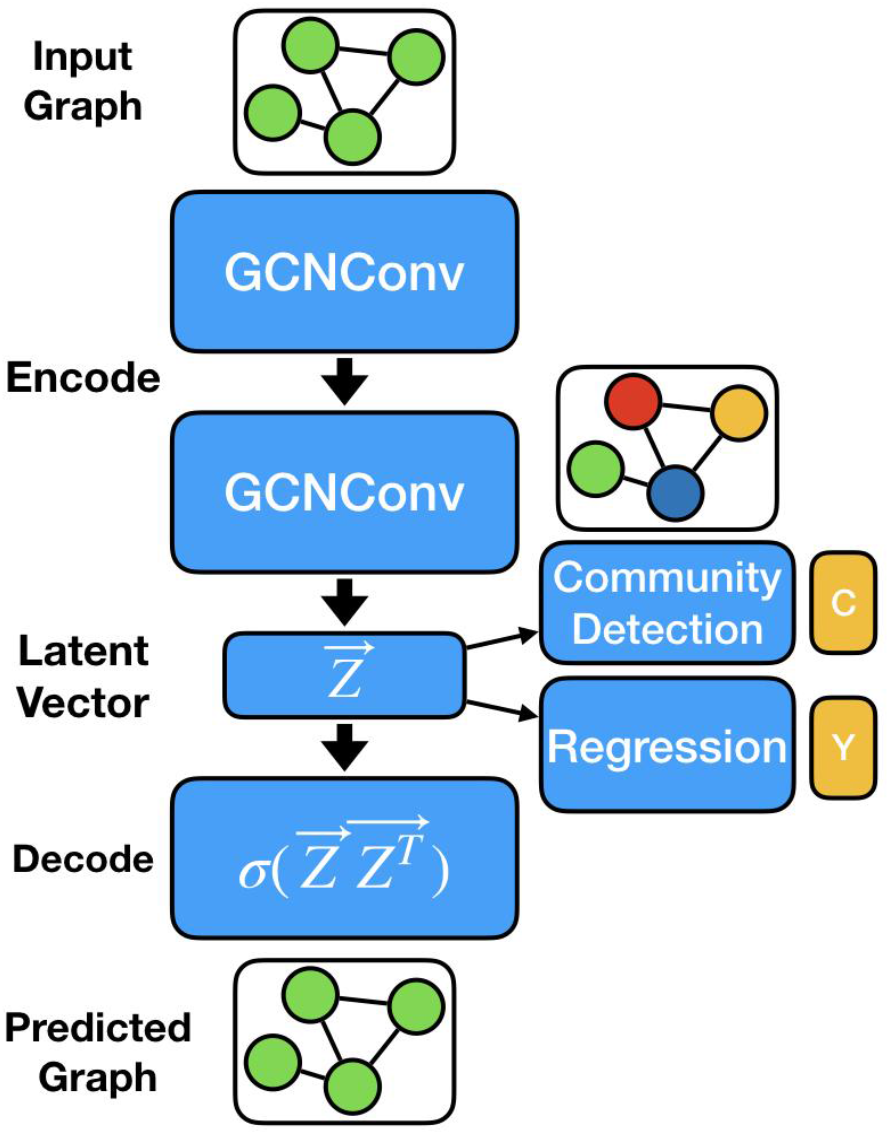
Illustration of GCN4R framework; graph, with adjacency matrix A and attribute matrix X are input into model; covariate information X is updated using successive applications of GCNConv operators (encoder) to yield lower dimensional representations of the nodes; latent vectors may be applied for community detection and regression tasks, or pairwise similarities may be computed to yield a predicted graph

**Figure 2:**
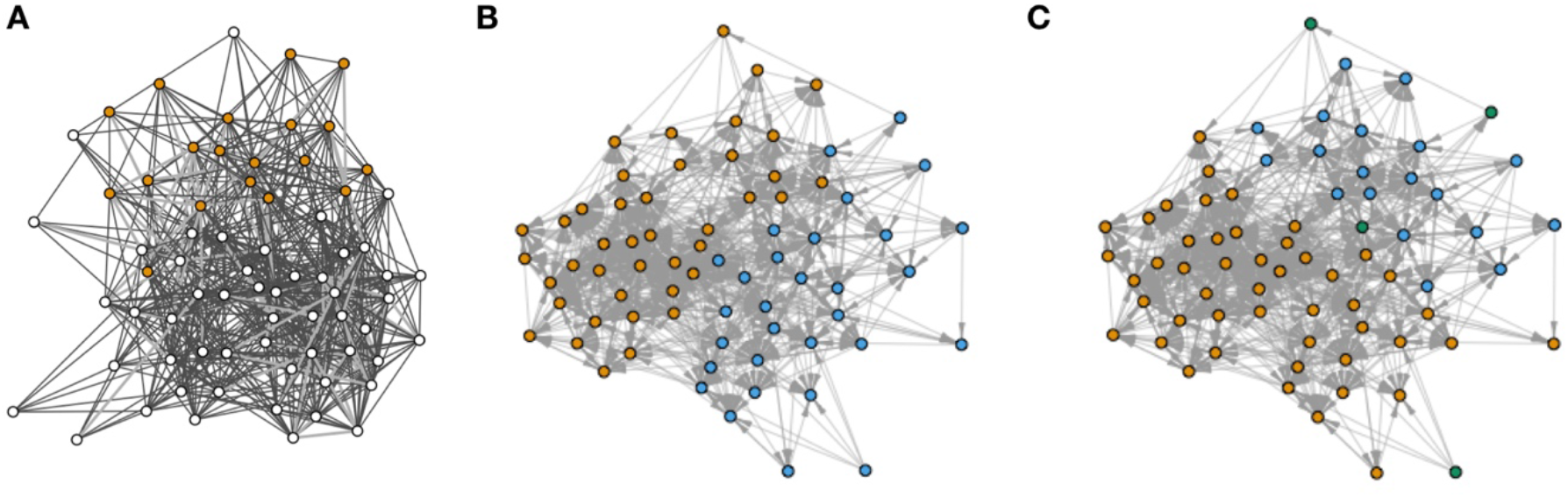
Visualization of the Lazega friendship network using the GCN4R platform. (A) Edges indicate ties between actors. Nodes colored in white based on if the lawyer works in the second law firm (office2 variable from design matrix). (B) Node indicative of lawyer practice and (C) node indicative of lawyer office

**Figure 3:**
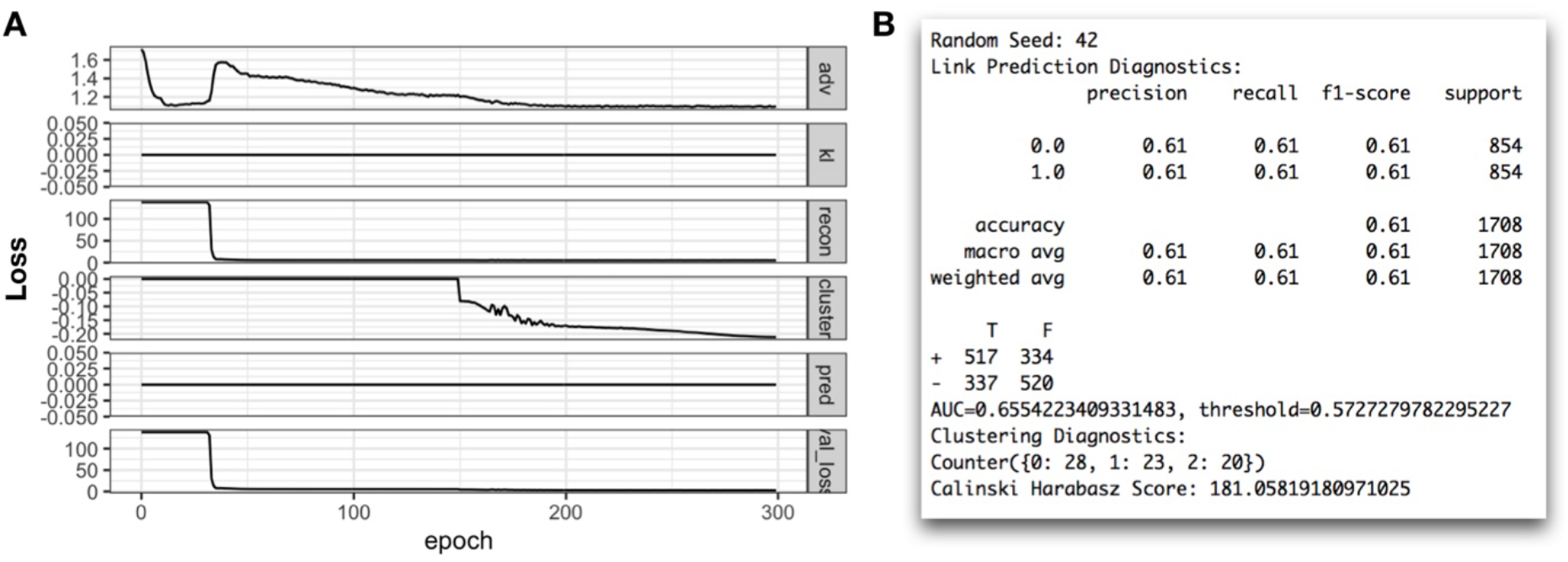
A) Diagnostic plot demonstrating behavior of various loss functions during fitting; B) Link Prediction and Community Detection Results as Output from the Summary command

The diagnostic plot displays components of the model objective, plotted over the training iterations. Of note in Figure 3A are the adversarial losses, reconstruction losses, cluster and validation losses, while the KL and prediction (classification/regression), losses have been removed by setting their weights *λ* to 0. We hope to see that all losses are decreasing over training iterations, indicating that the model is training towards convergence. The user should also check the scale of each of the loss components. If one of the components of the evaluated objective is of far greater magnitude than the others, then it is possible for the task being optimized to dominate the objective function and prohibit the successful optimization of other tasks. For instance, if the reconstruction loss is too large compared to the cluster loss, it is possible to fail to detect communities. Nonetheless, establishing the proper balance between the losses through coarse experimentation can lead to powerful regularization of the model objective for improved optimization.

The results of a cluster fit are stored in an R list extracted via the extract.results function and contain information such as the original and predicted graphs (extract.graphs), and the cluster assignments for each node (extract.cluster). The GCN4R package provides additional subroutines to extract measures of model fit in addition to the aforementioned diagnostics of the display of the model’s loss via the *summary* command (Figure 3B).

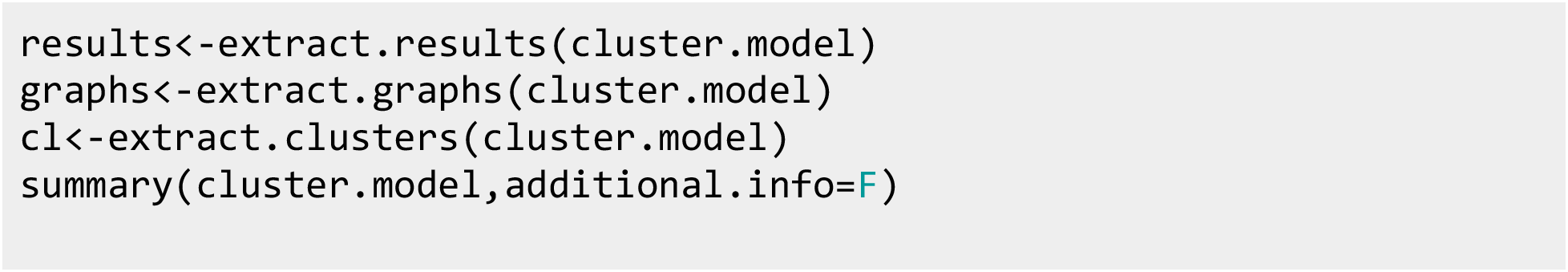

Here, a report was generated that details the overall accuracy for reconstruction of the network via the decoder mechanism (*p*_θ_(*G*|*Z*) = σ(*ZZ^T^*)) as described in the methods. The decoder outputs a probability score for the presence of a possible edge. The graph may be made more complete/sensitive by increasing this threshold score, at the expense of being less specific for the prediction of edges (capturing more false positives and fewer true negatives). An optimal threshold is calculated using Youden’s index to capture the maximum tradeoff between the sensitivity and specificity of link probability^44,45^. The area under the receiver operating curve (AUC) assesses whether a randomly selected edge which actually exists is assigned a higher predictive probability than a non-existent edge, which is also an overall measure of accuracy of the link prediction across a wide range of thresholds. Also displayed above is a summary of the number of individuals assigned to a cluster. Since each model fit is assigned a latent distribution of actors, within-cluster dispersion of actors can be assessed using the Calinski-Harabaz (CH) metric^46^, which is the ratio between the within-cluster dispersion and the between-cluster dispersion. As this score decreases, there is less mixing between the clusters and a lower potential for link formation across clusters since edge formation is based on the latent positions of the actors. The clustering and link prediction metrics may be compared between fitted models as an assessment of model fit.

### Model Visualization

Finally, the original and predicted networks of the cluster GNN model may be displayed using plot command as shown in Figure 4:

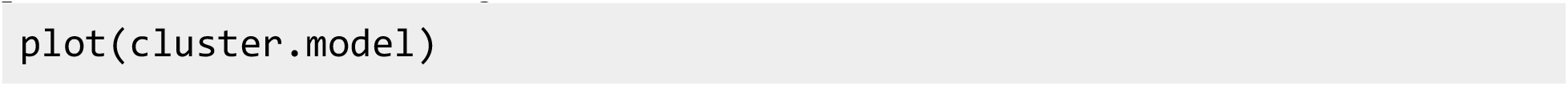

**Figure 4:**
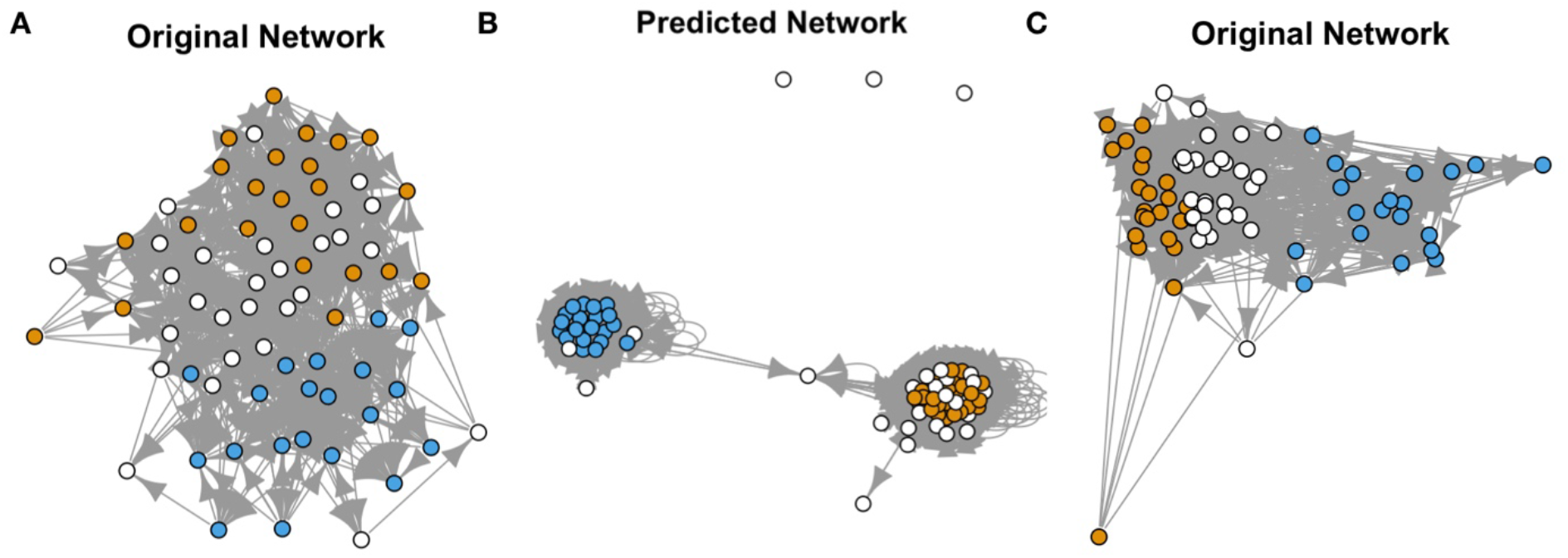
Visualization of Network After Model Fitting; Nodes are Colored by Cluster Assignment for: A) original network; B) predicted network (note how connectivity between clusters is diminished); C) original network ties plotted with neural network embeddings

These images may be saved to a file using the following command:

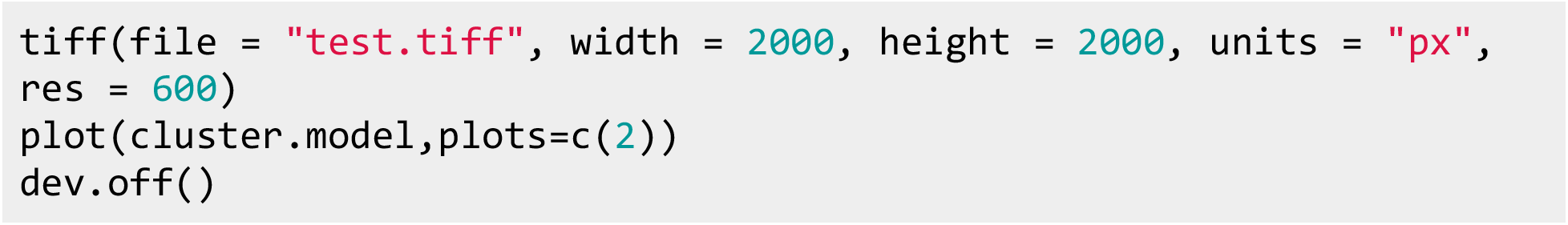

If one wishes to plot the learned latent positions of the actors, they may do so by adding the latent=T flag to the plot function:

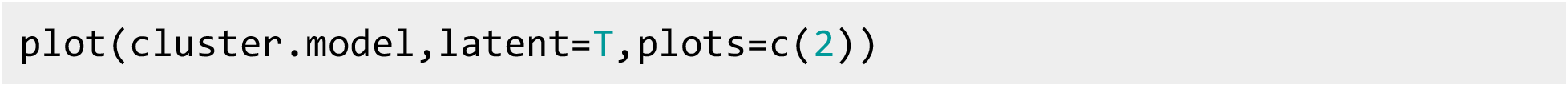

## Demonstration of Potentially Useful GNN Model Outputs

### Plotting Attention Matrices

We plot the attention mechanisms (weighted unipartite graph *α_i,j_*) for each layer of the cluster model using the following command:

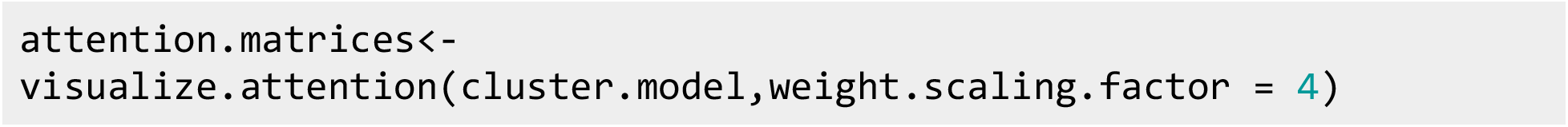

Here, we see that layer 1 (Figure 5A-B) tends to have sparse attention weights between peers, while layer 2 of the network (Figure 5C-D) is highly interconnected. Note how lawyers that are located on the periphery of the network diagram appear to have higher attention with the peers of their respective cluster. Given that members of formed communities should have similar connections and attributes, the attention weights visualized in Figure 5 may reflect the efforts of the model to “pull” lawyers that appear spuriously connected towards a community, perhaps based on factors that are outside dyadic independence.

**Figure 5:**
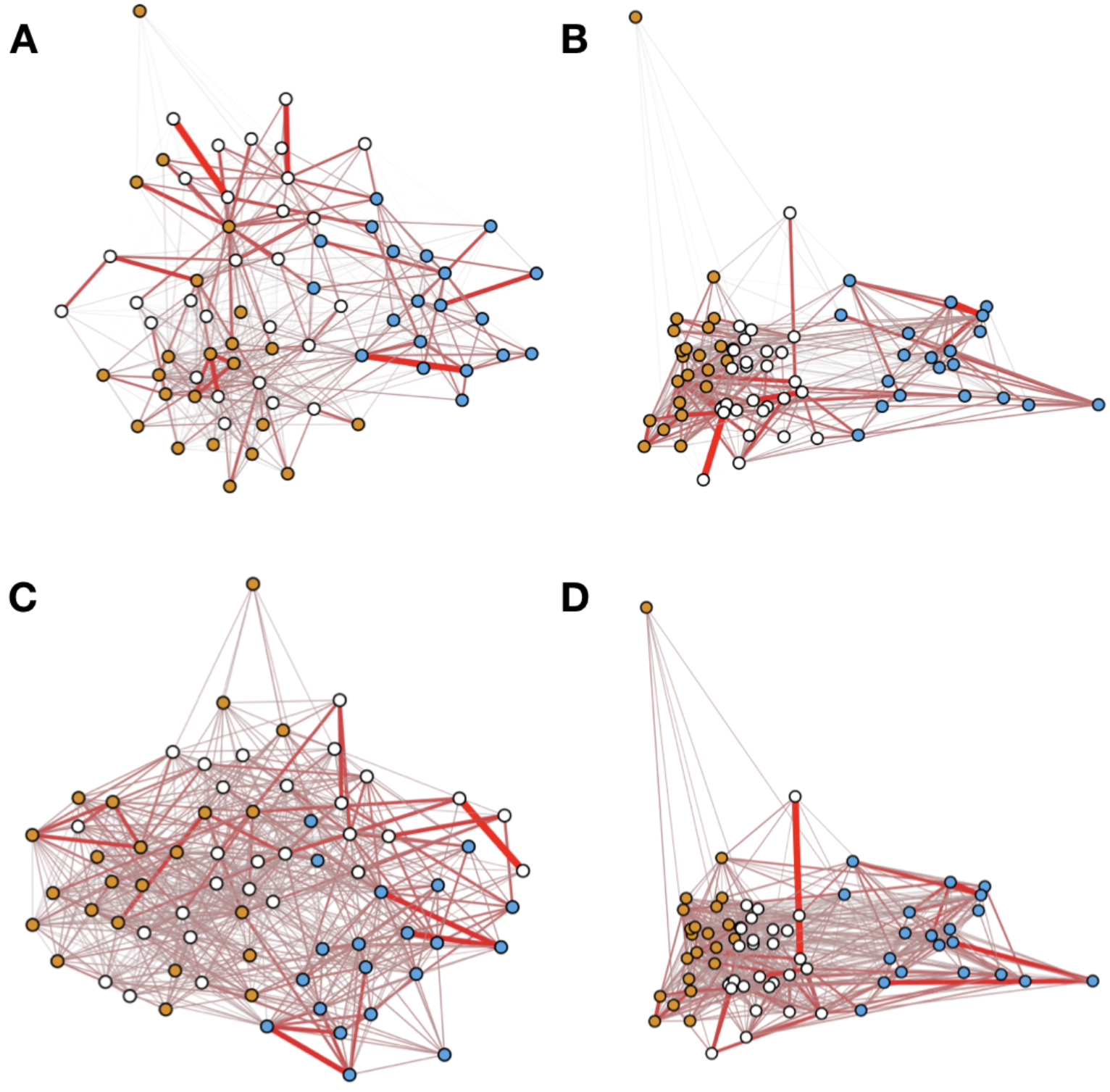
Visualization of Attention Weights; darker, thick red indicates higher weight between node and neighbor; color of node indicates cluster membership for: A,B) GATConv1; C,D) GATConv2; B,D) Plotted using uncovered neural network embedding layout

### Visualizing Integrated Gradients

Via post-hoc processing we are able to estimate which predictors are important for a given cluster assignment:

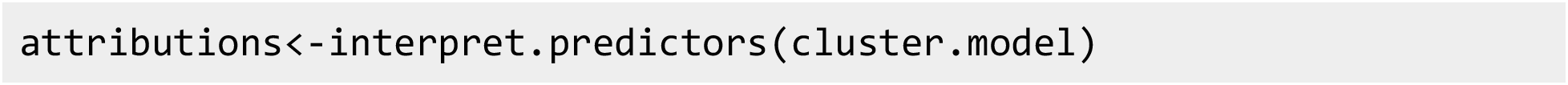

Figure 6A shows that age, seniority, and the lawyer’s law firm of practice were important contributing factors for selection into cluster “0”, though we do not present in this study the effect of predictor normalization on the importance of a particular variable. The importance of age and seniority for the formation of friendship clusters has been documented in prior studies on the Lazega network for community detection^47,48^.

**Figure 6:**
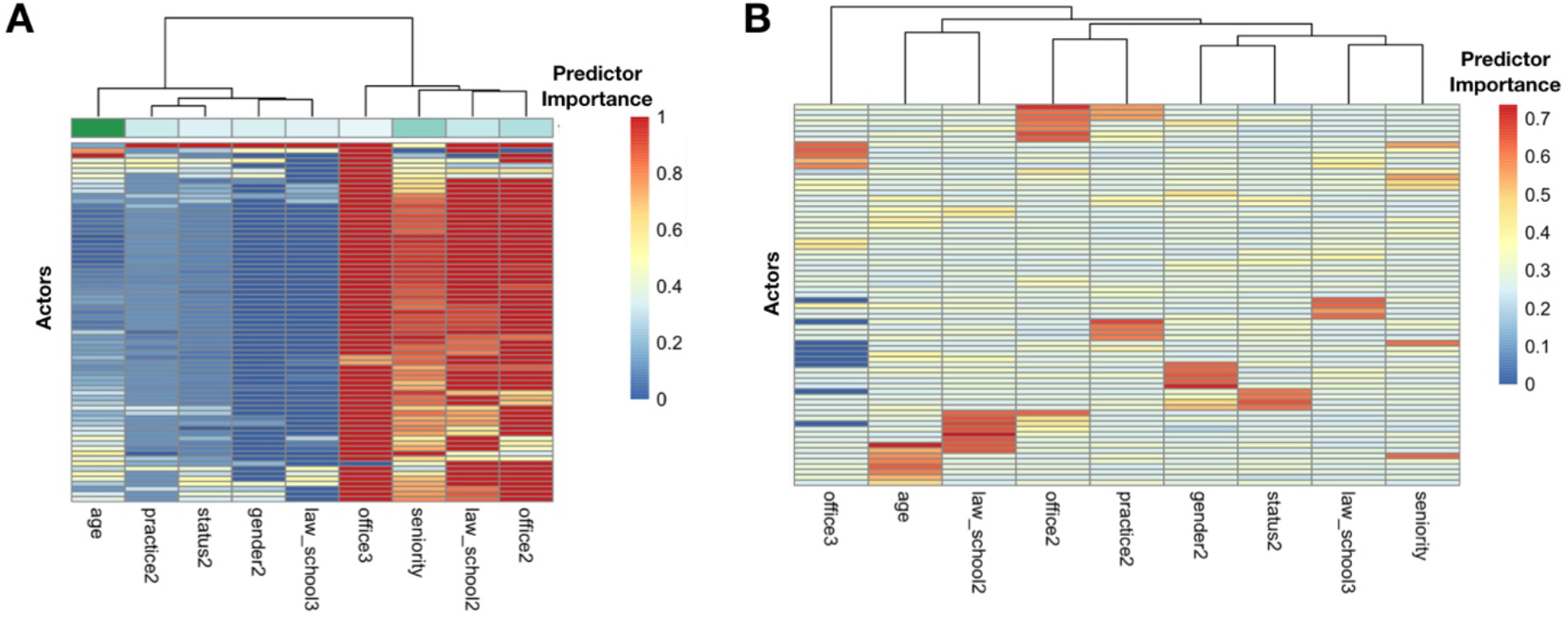
Visualization of Node Level Attributions of Predictors using: A) Integrated Gradients; red value indicates more important feature with respect to cluster 0 membership assignment; blue indicates low importance; importances have been summed column-wise to yield green color track on top; darker green indicates more important predictor; B) Predictor Masking via GNNExplainer; red indicates important predictor for the assigment of that cluster for the particular node; the left tracking bar indicates cluster membership for each node; actors were ordered separately for parts A) and B) via hierarchical clustering

### Egocentric Subgraph Extraction

Here, we refit the GNN using the GCNConv backbone:

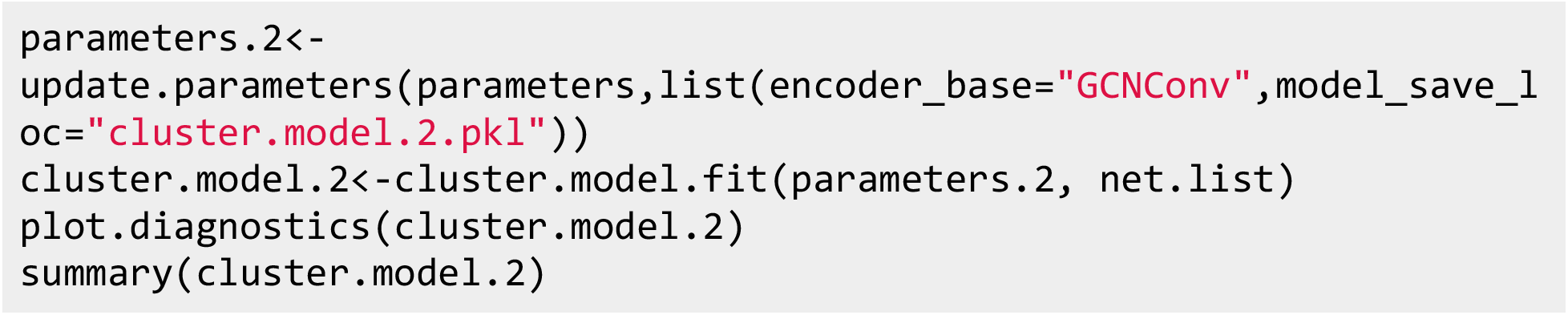

Results from the GNNExplainer can be returned using the following command:

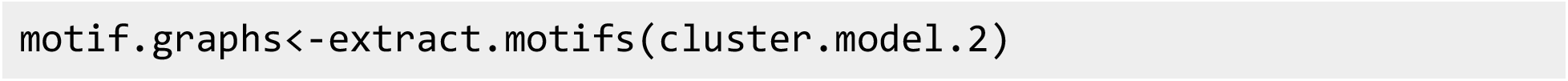

The predictor mask, *F*, that indicates the predictor importances for each node is returned with (Figure 6B):

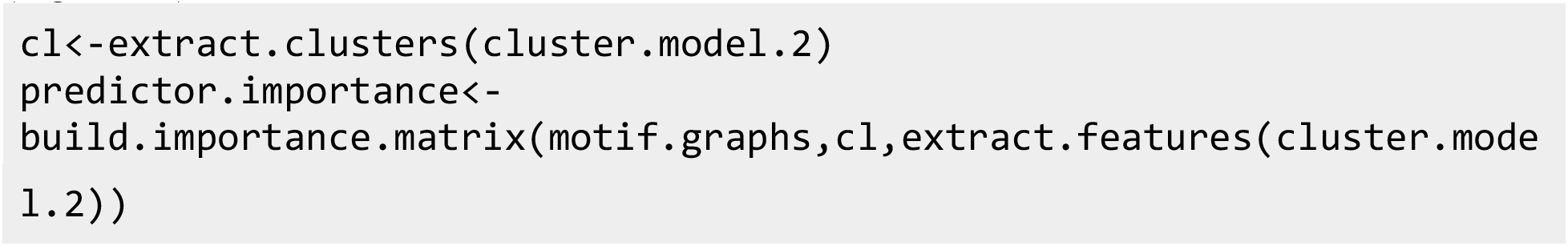

Interrogating the learned subgraph *M* of lawyer 60, pruning ties with an importance score less than 0.1 (Figure 7A):

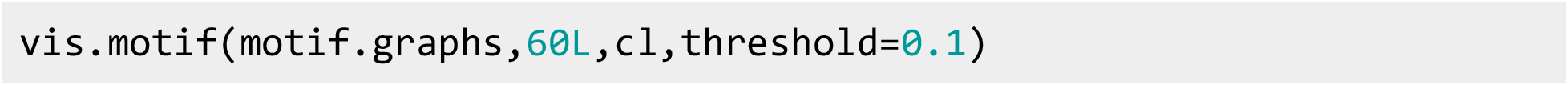

**Figure 7:**
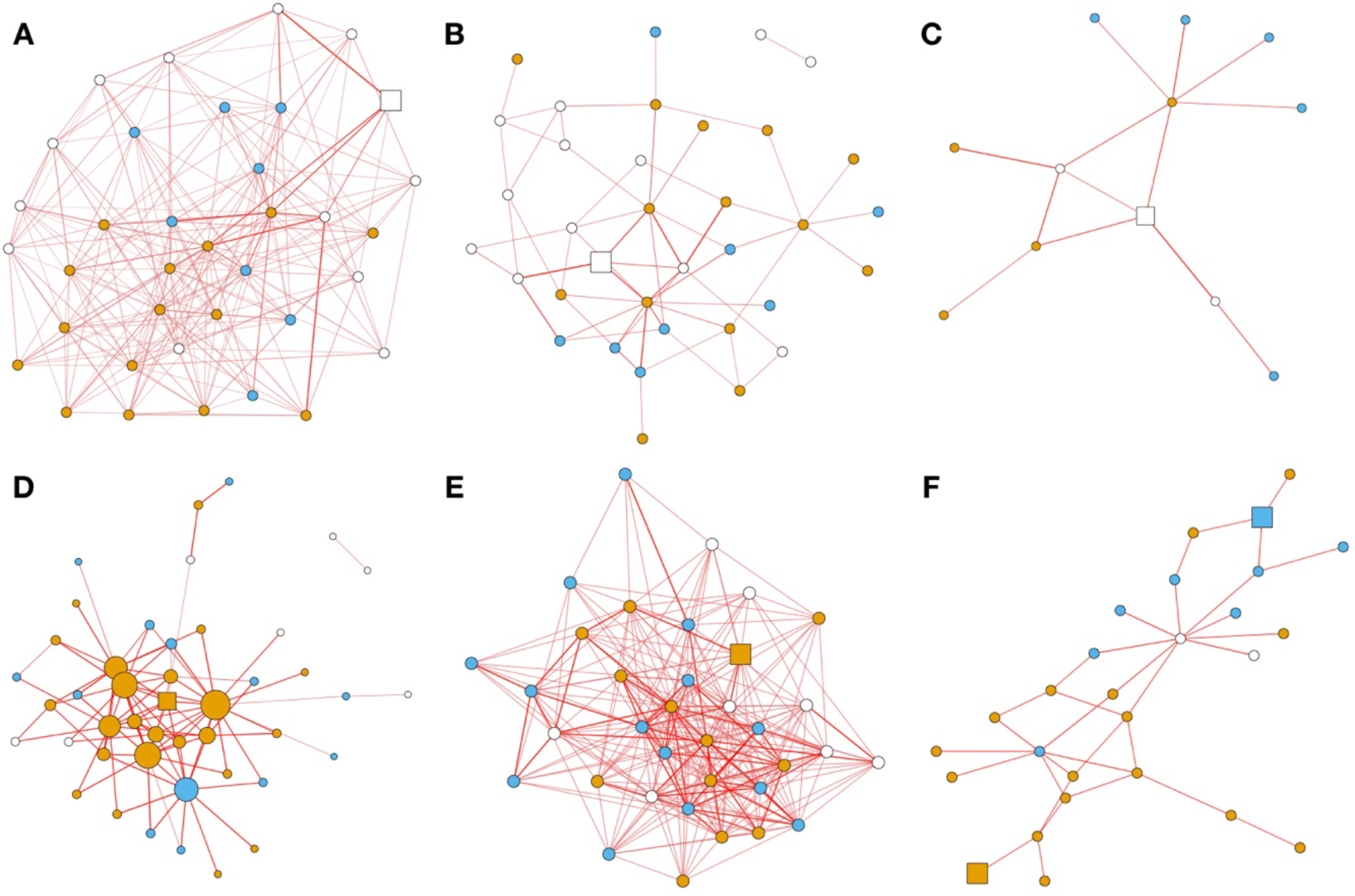
Visualization of extracted egocentric subgraphs (subgraphs of observed k-step subgraphs pruned for important ties) extracted using GNNExplainer (vis.motif); all squares indicate the node that is being interrogated; nodes colored by cluster assignment; edges thickness and darkness proportional to edge importance: A) subgraph that is most predictive of node 60’s cluster (white square) assignment given small threshold to include more higher order ties; B) increasing the importance threshold to 0.3 centers white square and lower order relationships are considered; C) further increase in importance threshold yields important ties within 2-path; D) subgraph for predicting lawyer 23’s cluster assignment (orange square); node size related to strength centrality measure of nodes; E) complete 2-step neighborhood around lawyer 23; F) consideration of integrating multiple nodes’ (blue and orange squares) learned subgraphs

The white square indicates that the contributing subgraph for node 60 should be inspected. Thicker and darker edges indicate more important connections by merit of maximizing the information needed to assign the correct cluster. These important connections are not always in the immediate vicinity of the node under inspection and in fact may represent second or third order connections. While this large subgraph demonstrates instances of community separation, we can prune it further by increasing the threshold to focus on these more important connections, thereby highlighting more specific motifs to focus on (Figure 7B-C):

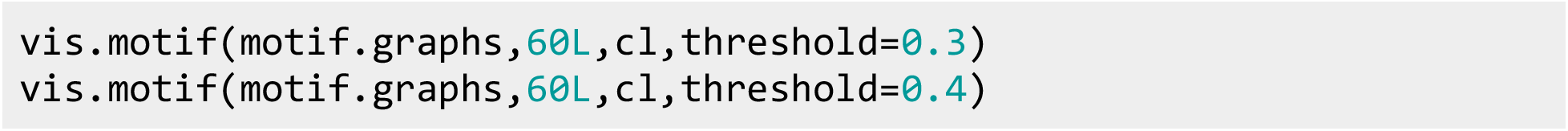

In Figures 7B-C, it would appear that some of the more important nodes that lawyer 60 is connected to are related to lawyers of a different cluster and define sharp boundaries. In another case, we interrogated the lawyers associated with node 23 and sized the nodes according to the strength centrality measure, which is similar to a degree measure, but instead sums the weights (learned by the explainer) of the connected edges (Figure 7D):

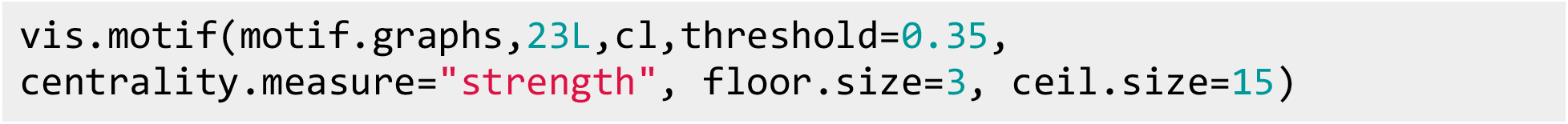

Except for two nodes, the geodesic distance between lawyer 23 and all of the other nodes (n=71) is less than or equal to two. Therefore, the egocentric network of lawyer 23 is contained within a 2-step neighborhood and most of the entire graph may be described by this neighborhood (Figure 7E). If we treat the 2-step neighborhood of lawyer 23 as the network, all nodes are retained and accessible within 2 ties. However, if we utilize the ego-centric network as learned by the GNNExplainer, 71% of the remaining edges in the network (237 out of 333 edges; 96 edges remain; original network has 854 edges) were pruned from this 2-step neighborhood (masked out; Figure 7D). Furthermore, the GNN explainer pruned some of the edges that are within 1-step, subsequently causing the geodesic distance between lawyer 23 and two other lawyers of the network to increase to 3. Although the distance between the lawyers have increased, the analysis has revealed specific and relevant information pathways (i.e., potentially more relevant transitive ties with higher order clustering) pertaining to the prediction as opposed to less but perhaps more immediate relationships. We have included an illustration of this concept in Figure 8.

**Figure 8:**
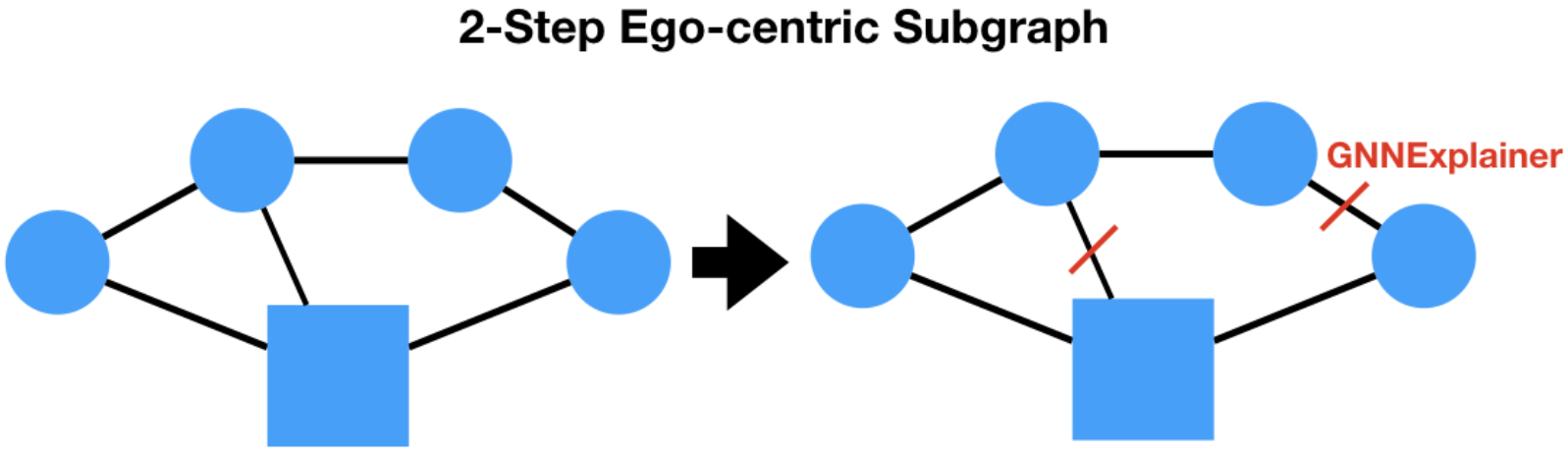
Illustration of GNNExplainer pruning ties from 2-step neighborhood of square node

In Figure 7F, we see that lawyer 23’s cluster assignment is heavily influenced by the assignment neighbors that are strongly connected themselves. One may also specify multiple nodes to assess for motifs (i.e., combining the subgraphs of both explained nodes), where all explained nodes are denoted using the square symbols (Figure 7F):

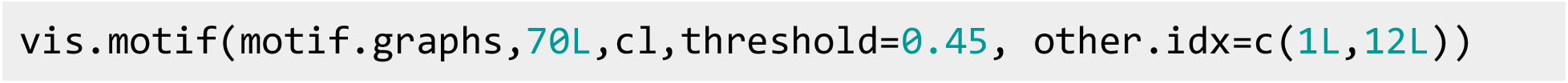

Clearly, some of the derived egocentric subgraphs could potentially contain spuriously assigned edges. Recall the methodology from section “**Perturbation Methods for Assessing Significance of Important Predictors, Edges and Motifs**”. In Figure 9, we demonstrate the application of this methodology pertaining to lawyers 37 (Figure 9A-B) and 61 (Figure 9C-D). We perturbed edges of the input graph with *p_pos_* = 0.3 and *p_neg_* = 0.05 across 180 bootstrapped networks to assess and remove statistically insignificant edges from egocentric subgraphs and plotted the original and resulting subgraphs using the following code:

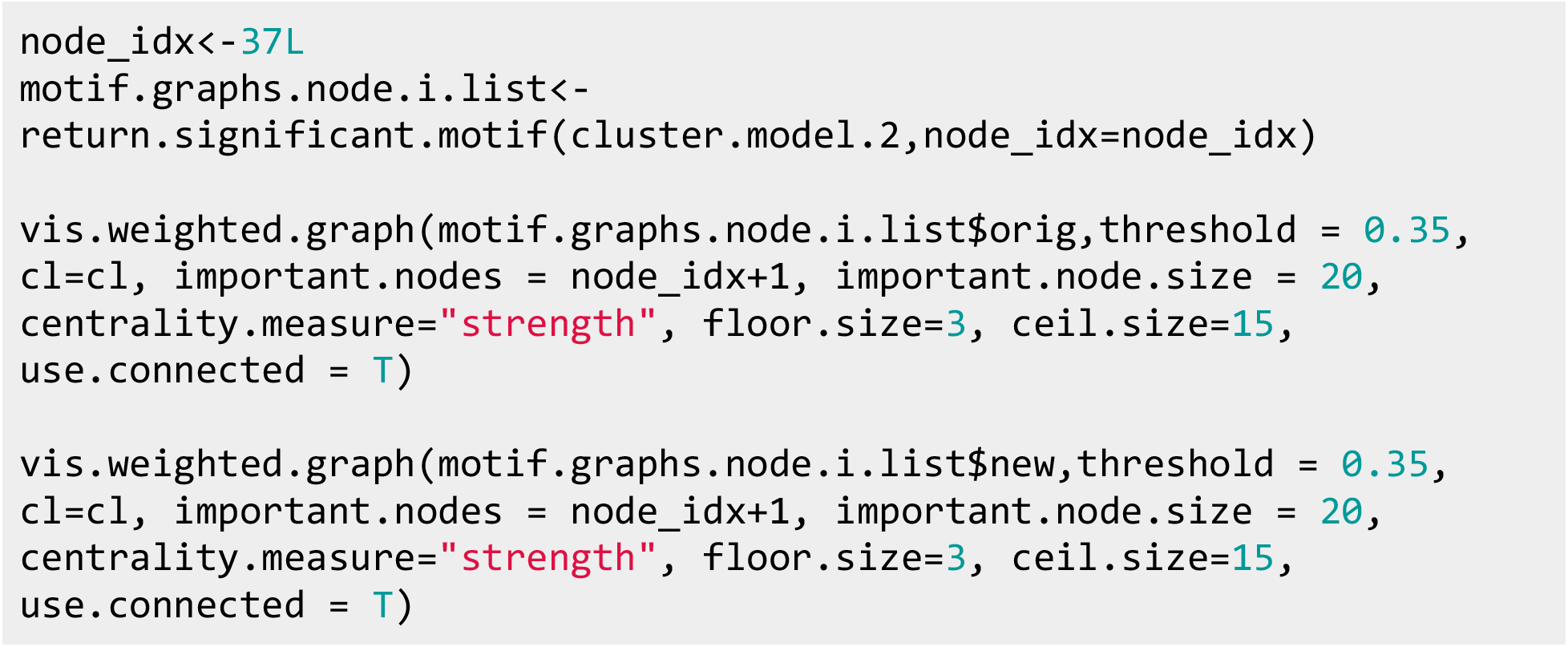

**Figure 9:**
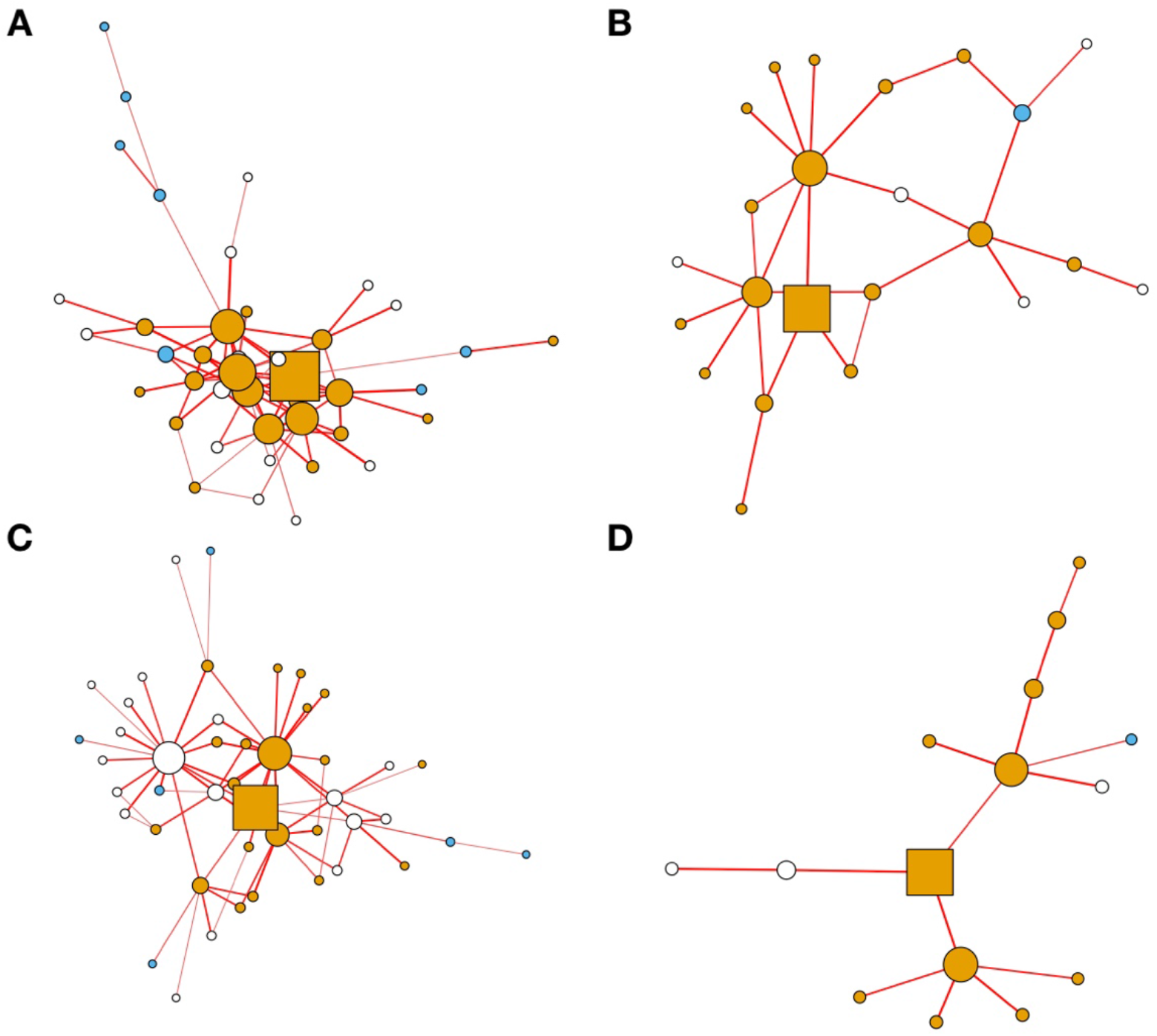
Visualization of extracted egocentric subgraphs (subgraphs of observed k-step subgraphs pruned for important ties) extracted using GNNExplainer (vis.weighted.graph); all squares indicate the node that is being interrogated; nodes colored by cluster assignment; edge thickness and darkness proportional to edge importance. Visualization for lawyers: A-B) 37 and C-D) 61. The original egocentric subgraphs are depicted in A,C) after setting the threshold to 0.35. Application of the one-sample Wilcoxon signed rank test which compares edge statistics to that of 180 perturbed graphs reveals simpler information pathways in B,D) under the same 0.35 cutoff threshold, where potentially spurious edges have been filtered after Bonferroni correction

The interpretation of the egocentric subgraphs extracted for lawyers 37 and 61 (Figure 9B,D) are as follows:

- Lawyer 37 (Figure 9B) reach extends largely to three influential lawyers (three larger orange circles), each of whom have influenced lawyer 37’s community assignment to the same cluster. Each of these three lawyers are likely in communication with one another, either directly or indirectly through transitive relationships.
- Lawyer 61 (Figure 9D), in contrast, is directly connected to two lawyers (two larger orange circles) who do not communicate amongst each other. However, these lawyers are well connected with other lawyers from the same community, who in turn indirectly influence the membership of lawyer 61.

While derivation of the egocentric subgraphs is important, using perturbation to assess for statistically meaningful subgraphs can help simplify complex ego networks in the context of the GNN prediction task.

### Node Importance

Although there are many other characterizations of important/influential nodes in a model ^29,49^, we have supplied a few heuristics from which to derive importance scores for these nodes.

Performance/disruptibility based node importance measures estimate the importance of a node by removing the node and its immediate connections (i.e. disrupting the network) and recording the change/decrease in performance of the model (e.g., node-level C-statistic for classification tasks, *R*^2^ for regression tasks, etc.). A node has greater importance if its removal would cause greater disruption. For clustering models, this is actualized as the change in cluster assignment when the node is removed, as measured through the V-Measure^50^, which is a normalized measure of mutual information (NMI) between the previous and perturbed cluster assignment (Figure 10A):

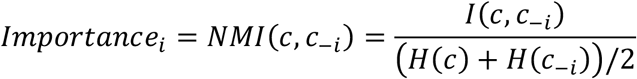

**Figure 10:**
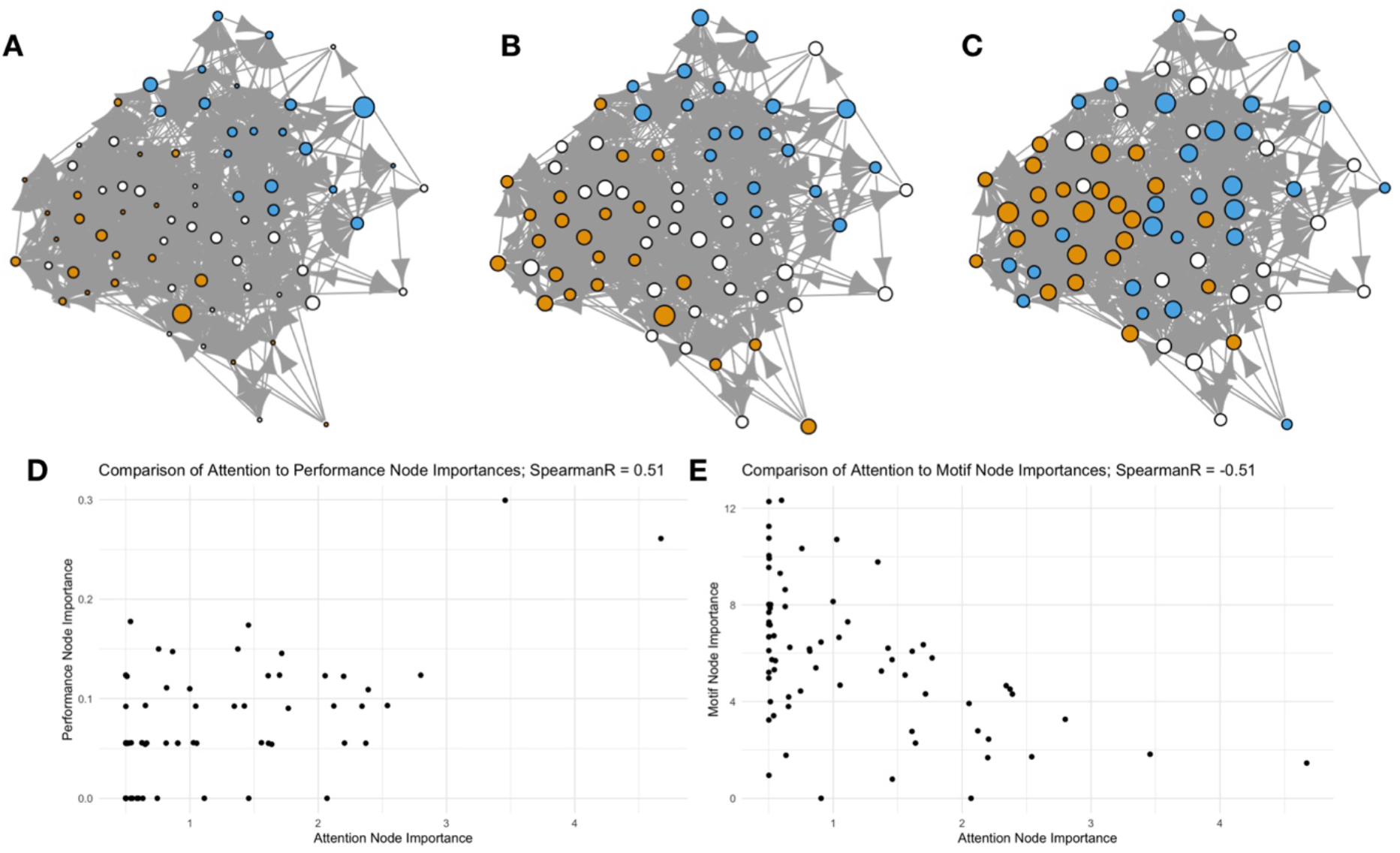
Visualization of Node Importance Measures over Original Lazega Network; nodes colored by cluster membership; node size is proportional to: A) performance importance; B) attention importance; C) motif importance; D) comparison between attention and performance importance; E) comparison between attention to motif importance

Here *I* is the mutual information between the original cluster assignments *c* and the perturbed cluster assignments *c_−i_* after removing node *i*, which has been normalized by the arithmetic mean of their respective entropies. Alternatively, for classification and regression tasks, one may calculate reductions in accuracy, C-statistics (area under the receiver operating curve; AUC), coefficient of determination *R*^2^, or F1-score per removal of the node; for instance, for node *i*:

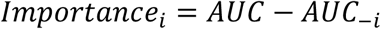

We have included code to calculate the *NMI* below:

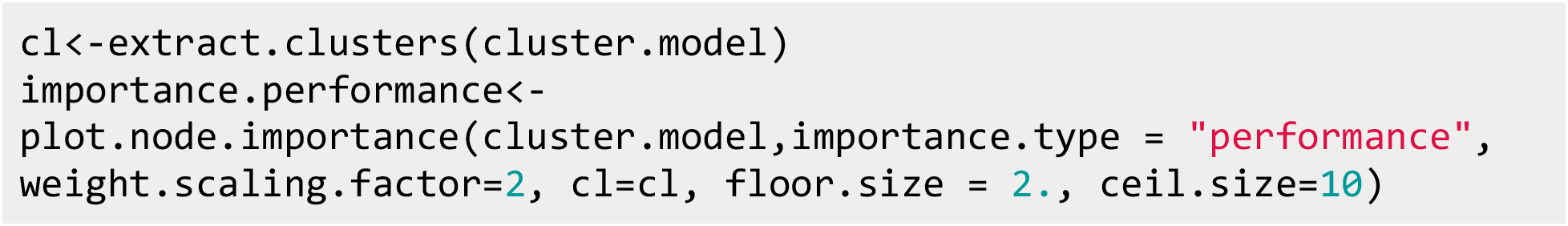

Gradient-based node importance sums the predictor-specific importance scores across node-level attributes, as derived using the integrated gradients algorithm.

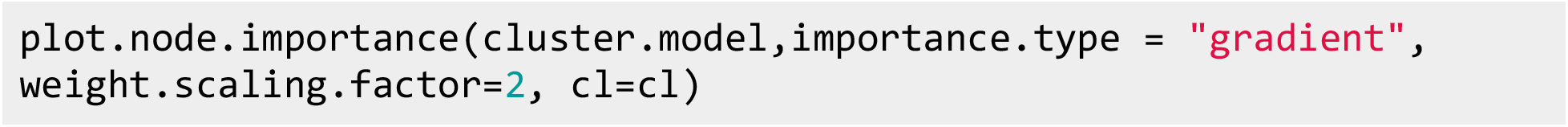

Attention based methods seek to derive an importance score for each node based on centrality measures that are weighted by the learned attention weights. These importance scores may be derived for each layer of the network and speak to how much information one node may receive from neighboring nodes (Figure 10B):

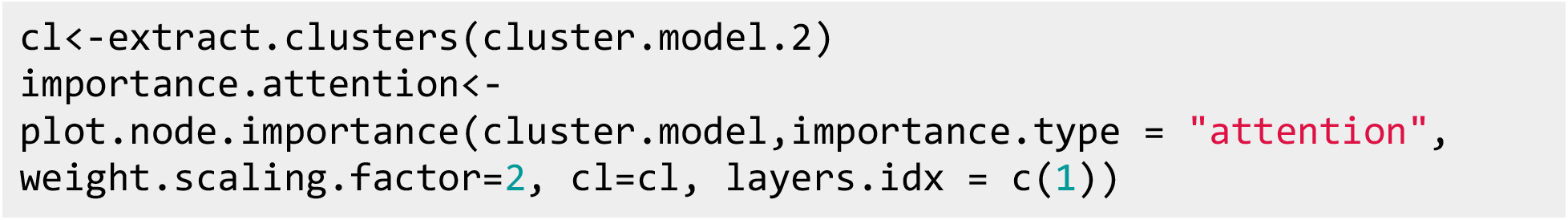

Motif-based methods compute the centrality of each node with respect to their learned subgraph mask (Figure 10C):

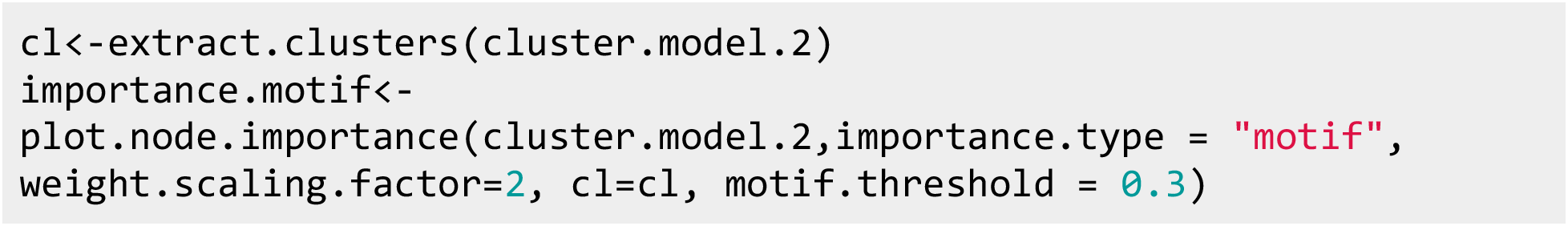

In this example, the performance and attention-based measures correlate positively with each other and the motif and attention-based measures correlate negatively with each other (Figure 10D-E). The significance/stability of the node-level importances can be assessed through perturbation methods outlined in section “**Perturbation Methods for Assessing Significance of Important Predictors, Edges and Motifs**”, though this has not been explicitly programmed.

### Alternative Modelling Optimization Schemes

Here, we increase this *λ_KL_* penalty from 0 for the prior fit models to 1e-2:

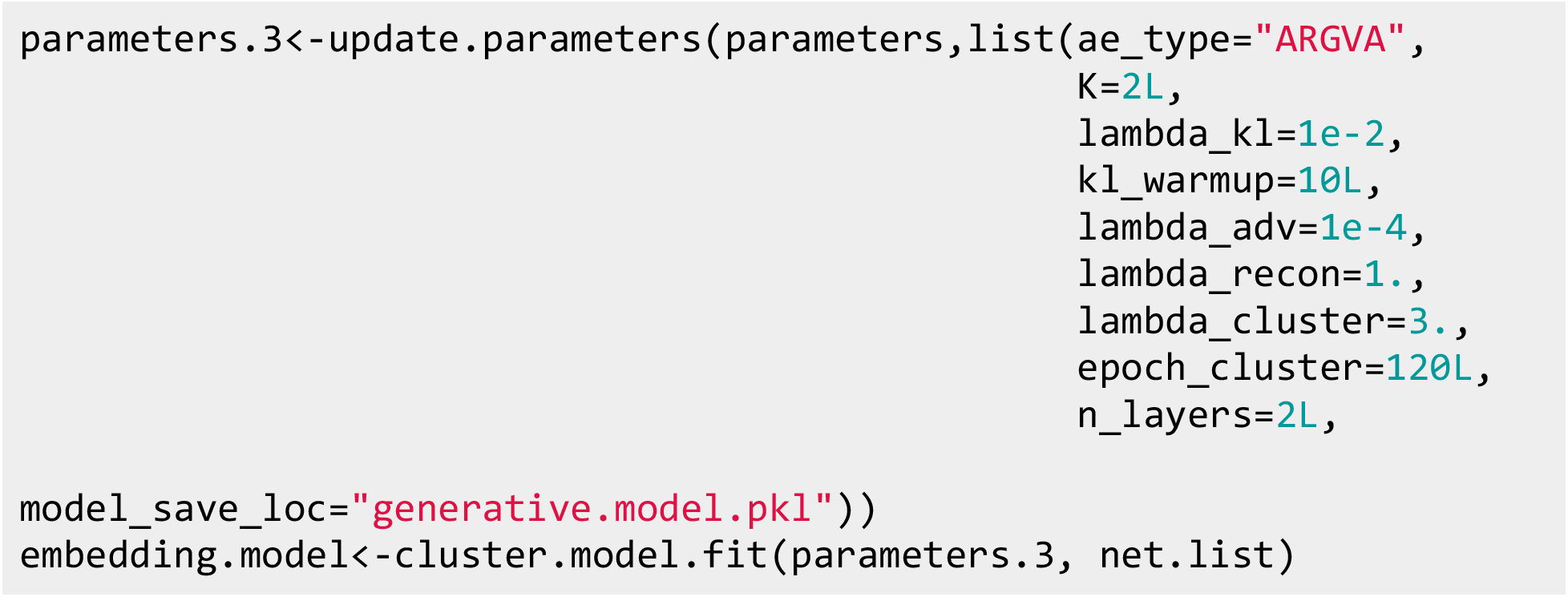

A preliminary check of the diagnostics indicates that the KL-Loss term is of the same magnitude as the reconstruction loss (Figure 11E); this indicates that there is heavy penalization of the latent actor positions towards multivariate normal. Near epoch 200, the clustering loss decreased towards its minimal value, which had allowed the fitting of two clusters. Inspection of the link prediction (Figure 11F) demonstrates that there were more false negative links predicted versus true positive, demonstrating lower sensitivity than random chance alone:

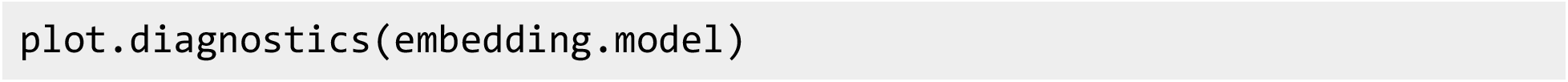

**Figure 11:**
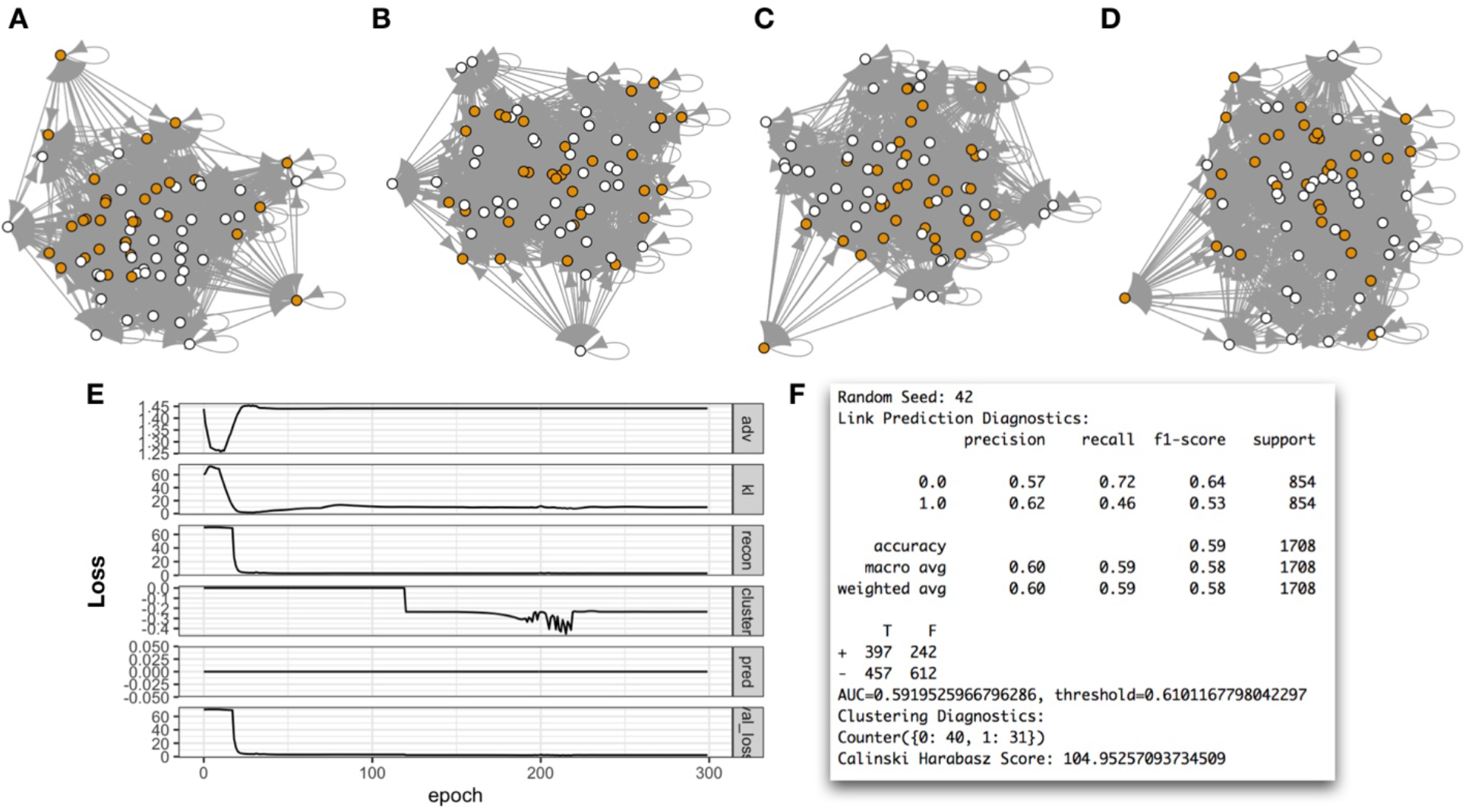
Visualization of Generated Networks; colored by learnt cluster assignments: A-D) four generated networks sampled from latent positions of actors; sampled latent positions have been plotted for each network; E) Diagnostic curves depicting each loss over time; F) link prediction and clustering statistics output from summary function

We perform 30 posterior draws from the latent distribution of actor positions of the lawyers of the network and plot four of the derived networks along with their latent positions:

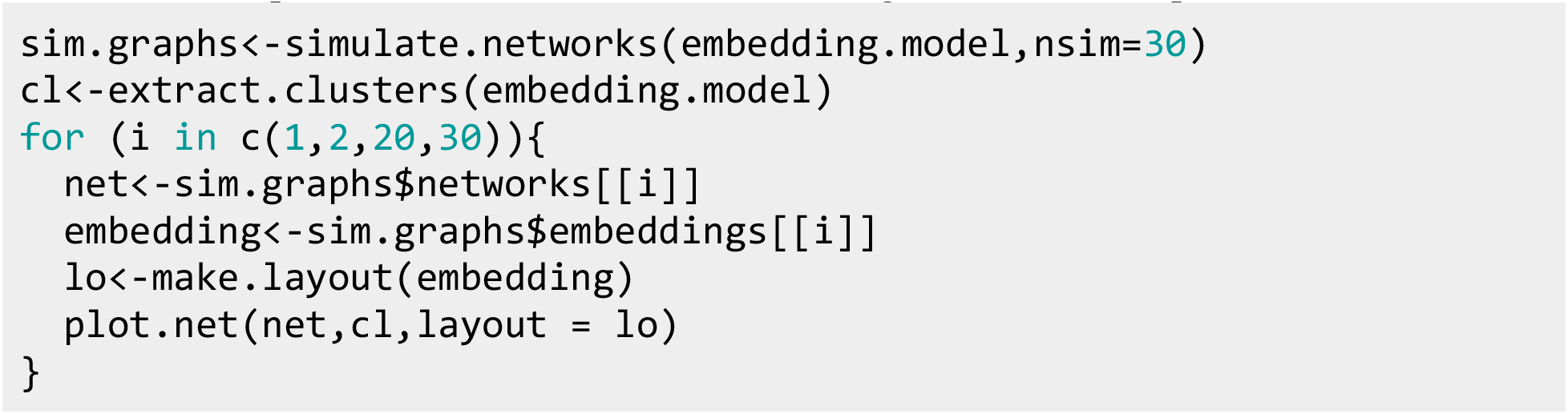

Each lawyer is colored by their original learned cluster assignments during the training of the model (Figure 11A-D). We see that there is rough cluster separation (CH from the generative model is less than that of the original cluster model), but sampling from the multivariate posterior latent distribution have yielded substantially different networks and latent positions.

### Classification and Regression

We now predict whether a lawyer was a part of law firm 2:

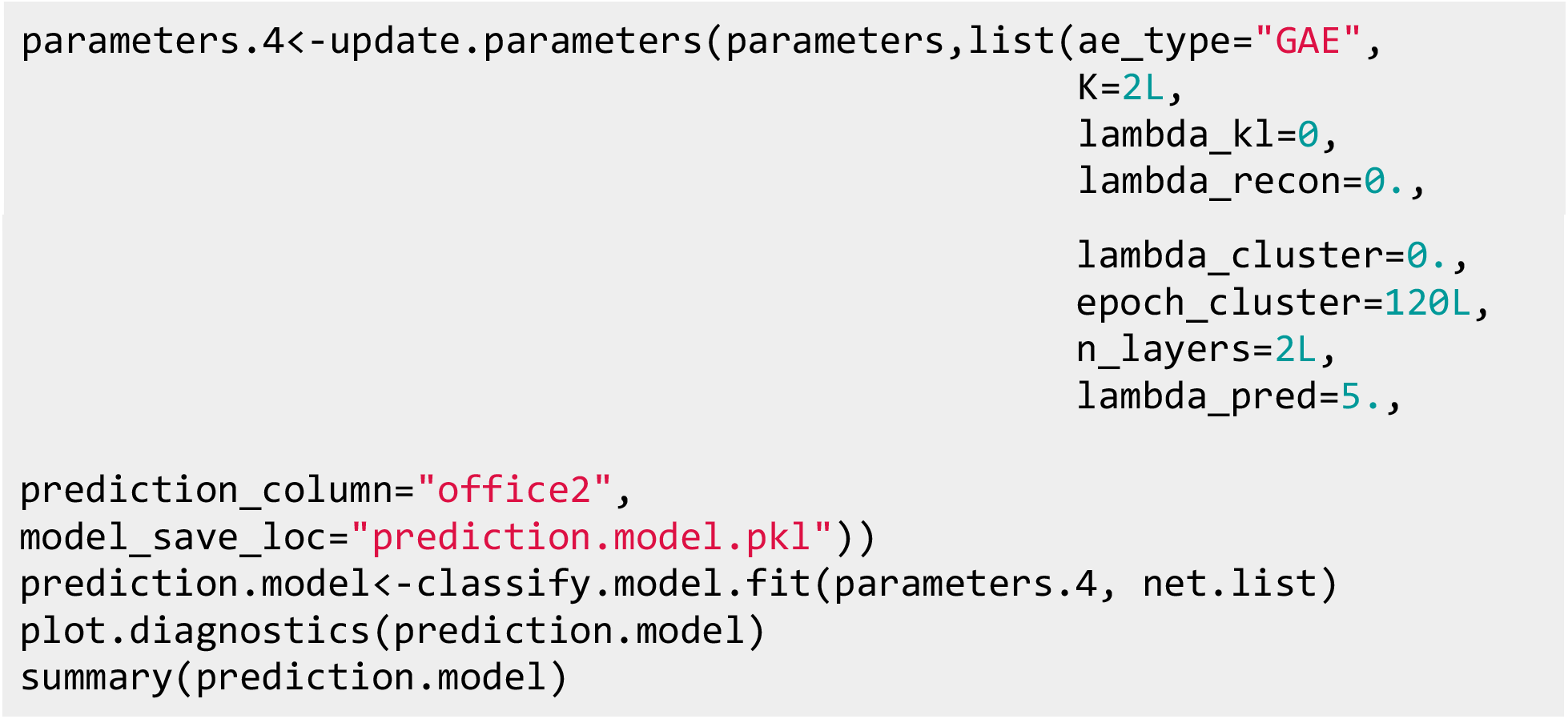

We focused purely on estimation of the lawyer’s office rather than other modeling objectives, though these other objectives may be introduced. We find that the model is able to perfectly classify which law firm each lawyer works at (Figure 12C). This is confirmed from the extraction of the latent vectors of the fit model (Figure 12A):

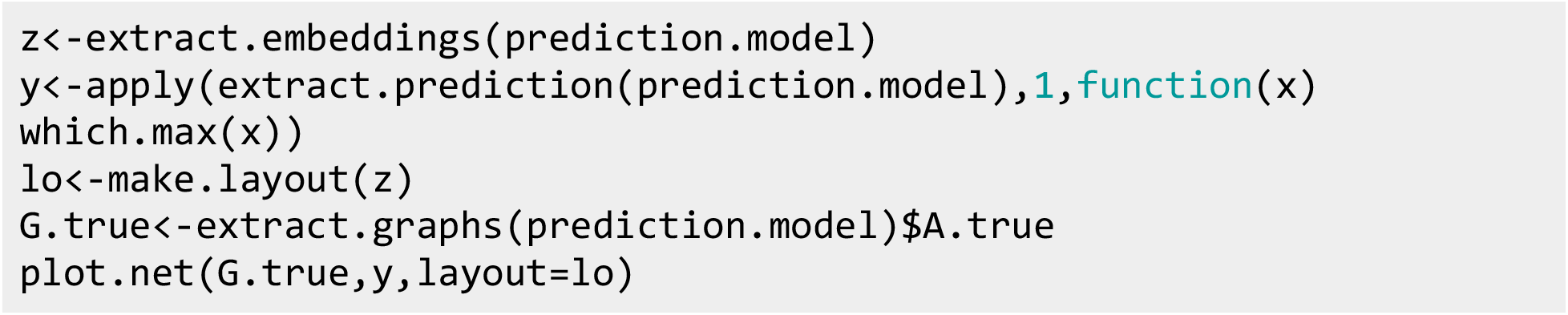

**Figure 12:**
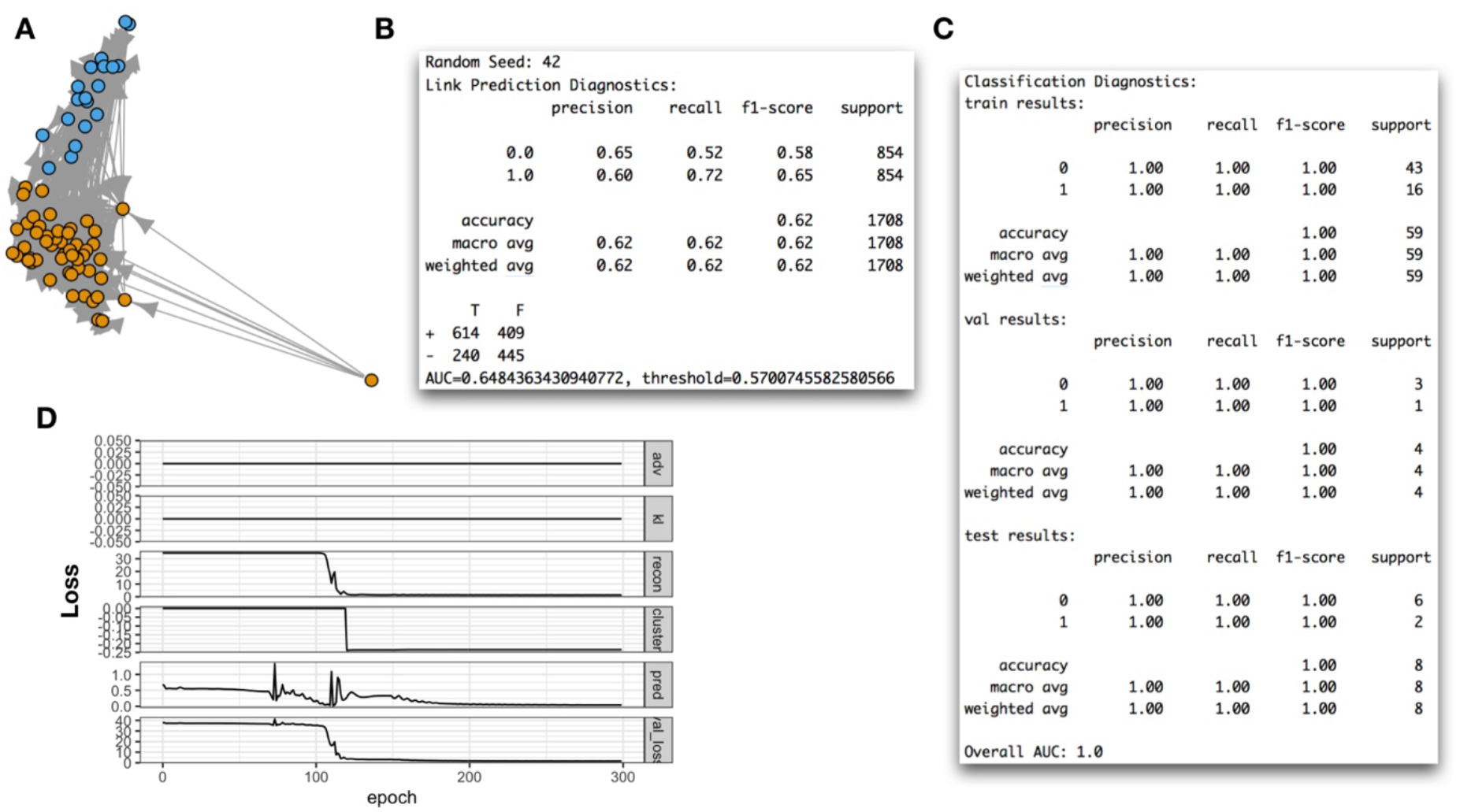
Classification results for whether lawyer worked at second law firm; A) latent positions of actors, colored by whether lawyer worked at second law firm; B) link prediction results via the summary function; C) classification results on smaller subsets of actors that constitute the training (n=59), validation (n=4) and test set (n=8) of actors; D) diagnostic loss curves plotted over training epochs

### Additional Comparisons to Traditional Approaches

The GNN methodology focuses more on prediction, while their statistical counterparts focus more on network understanding. Extensive comparisons of predictive performance are beyond the scope of this study^22,51^. However, in this section we present a few side-by-side comparisons.

As an example, the results from the cross-sectional network auto-correlation model (e.g., lnam) in estimating the social influence of actors on the behaviors of one another (for instance, which law firms the lawyers practice at), may be readily compared to the attention-based measures previously demonstrated. Given that one or more attention layers may be learned at a given layer, the matrix multiplication of these weights across layers may give an indication of which node has influenced the prediction of another within k-steps (for k GCN layers). We have included an example below (Figure 13), which compares the learned 4-step social influence on law firm selection for each lawyer as estimated by: 1) a cross-sectional network auto-correlation model, versus 2) compiling information extracted from four graph attention layers.

**Figure 13:**
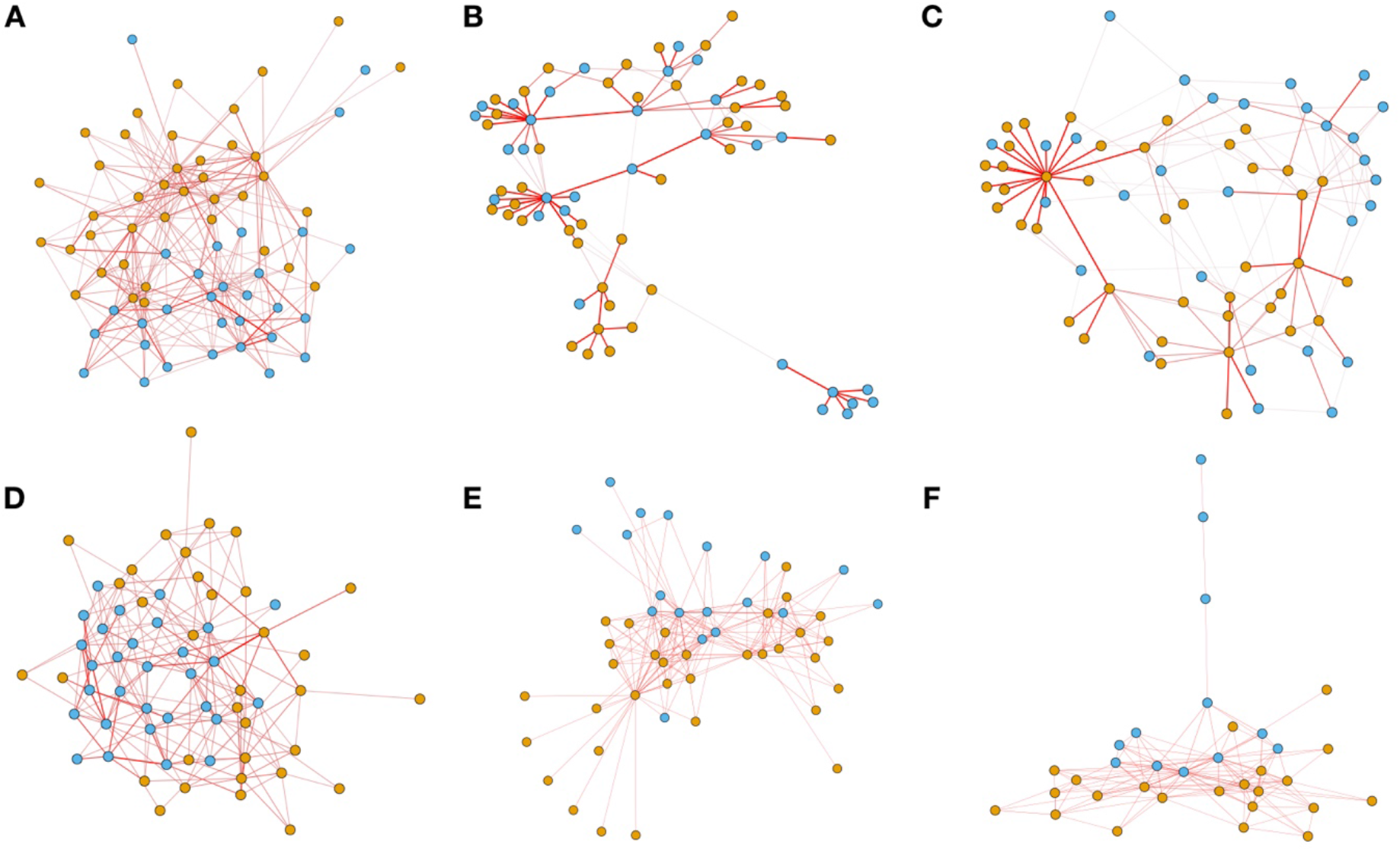
An Example of Social Influence Results from Prediction of Lawyer’s Law Firm (color of nodes in networks) from Seniority, Age, and Gender: Visualization of attention weights for GCN: A) Layer 1, B) Layer 2, C) Layer 3, D) Layer 4; these attention matrices are matrix multiplied together to form: E) the 4-step social influence network; this GCN social influence network is compared to: F) a social influence network derived after predicting law firm from a similar network autocorrelation model with MA(1) autoregressive dependency and inverse geodesic distance affinity

Figure 8 demonstrated the advantages of pruning ego-centric subgraphs for contextualizing the nodes of the network. As a final note, we visualize the differences that may be acquired from calculating Louvain modularity over the network, fitting a latentnet model, and clustering using the GCN, which incorporates the node-level predictors in its estimation (Figure 14).

**Figure 14:**
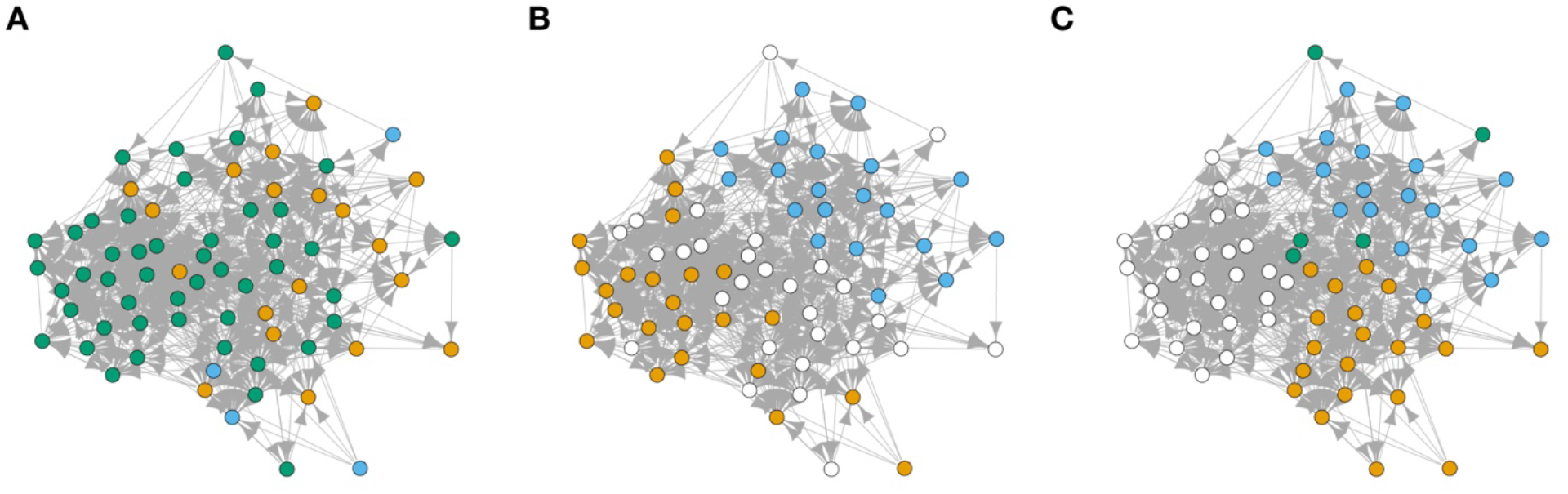
Comparison of community detection results after fitting a (nodes colored by communities): A) Bilinear Latent Space Model, B) Adversarially Regularized Graph Autoencoder with Spectral Clustering objective, C) Louvain’s modularity maximization algorithm

## Conclusion

In this tutorial paper, we provided a broad overview of possibilities of what can be accomplished with graph neural network approaches for learning on graph-structured data. Possible extensions to these methods would include the ability to generalize node embeddings across graphs (inductive learning), a comparison of multiple graphs by estimating graph-level embeddings (pooling mechanisms which learn the properties of the graph as a whole), predicting ties and categorial or continuous node/graph-level diffusion of status over time^52^, and software ported to R to improve visualization of large-scale networks using deep learning embeddings to summarize graphs via topological data analysis^53^. We emphasize additional opportunities to combine the insights that can be acquired from both statistical and machine learning techniques over networks. Graph Neural Network approaches for network analysis are coming of age and have helped researchers in the machine learning community gain a deeper understanding of actor positions in the network. As such, software solutions that simplify and extend these approaches to additional scientific communities (R users), are expected to facilitate additional research into the development and application of these methods for biological and social network analysis.

## References

1. O’Malley, A. J. The analysis of social network data: an exciting frontier for statisticians. Statistics in Medicine 32, 539–555 (2013).

2. Langfelder, P. & Horvath, S. WGCNA: an R package for weighted correlation network analysis. BMC Bioinformatics 9, 559 (2008).

3. van IJzendoorn, D. G. P., Glass, K., Quackenbush, J. & Kuijjer, M. L. PyPanda: a Python package for gene regulatory network reconstruction. Bioinformatics 32, 3363–3365 (2016).

4. Hevey, D. Network analysis: a brief overview and tutorial. Health Psychology and Behavioral Medicine 6, 301–328 (2018).

5. Delgado, F. M. & Gómez-Vela, F. Computational methods for Gene Regulatory Networks reconstruction and analysis: A review. Artificial Intelligence in Medicine 95, 133–145 (2019).

6. Handcock, M. S., Hunter, D. R., Butts, C. T., Goodreau, S. M. & Morris, M. statnet: Software Tools for the Representation, Visualization, Analysis and Simulation of Network Data. J Stat Softw 24, 1548–7660 (2008).

7. Handcock, M., Raftery, A. & Tantrum, J. Model-Based Clustering for Social Networks. Journal of the Royal Statistical Society: Series A (Statistics in Society) 170, 301–354 (2007).

8. Krivitsky, P. N. & Handcock, M. S. Fitting Position Latent Cluster Models for Social Networks with latentnet. J Stat Softw 24, (2008).

9. Goodreau, S. M., Handcock, M. S., Hunter, D. R., Butts, C. T. & Morris, M. A statnet Tutorial. Journal of Statistical Software 24, 1–26 (2008).

10. Blondel, V. D., Guillaume, J.-L., Lambiotte, R. & Lefebvre, E. Fast unfolding of communities in large networks. J. Stat. Mech. 2008, P10008 (2008).

11. Hoff, P. D., Raftery, A. E. & Handcock, M. S. Latent Space Approaches to Social Network Analysis. Journal of the American Statistical Association 97, 1090–1098 (2002).

12. Zitnik, M., Agrawal, M. & Leskovec, J. Modeling polypharmacy side effects with graph convolutional networks. Bioinformatics 34, i457–i466 (2018).

13. Tan, Q., Liu, N. & Hu, X. Deep Representation Learning for Social Network Analysis. Front. Big Data 2, (2019).

14. Nelson, W. et al. To Embed or Not: Network Embedding as a Paradigm in Computational Biology. Front. Genet. 10, (2019).

15. Cui, P., Wang, X., Pei, J. & Zhu, W. A Survey on Network Embedding. arXiv:1711.08752 [cs] (2017).

16. LeCun, Y., Bengio, Y. & Hinton, G. Deep learning. Nature 521, 436–444 (2015).

17. Zhu, J. et al. Generalizing Graph Neural Networks Beyond Homophily. arXiv:2006.11468 [cs, stat] (2020).

18. Zhou, J. et al. Graph Neural Networks: A Review of Methods and Applications. arXiv:1812.08434 [cs, stat] (2019).

19. Wu, Z. et al. A Comprehensive Survey on Graph Neural Networks. (2019).

20. Fey, M. & Lenssen, J. E. Fast Graph Representation Learning with PyTorch Geometric. arXiv:1903.02428 [cs, stat] (2019).

21. Zhang, M. & Chen, Y. Link Prediction Based on Graph Neural Networks. arXiv:1802.09691 [cs, stat] (2018).

22. Qiu, J. et al. DeepInf: Social Influence Prediction with Deep Learning. Proceedings of the 24th ACM SIGKDD International Conference on Knowledge Discovery & Data Mining 2110–2119 (2018) doi:10.1145/3219819.3220077.

23. Pan, S. et al. Adversarially Regularized Graph Autoencoder for Graph Embedding. arXiv:1802.04407 [cs, stat] (2019).

24. Lazega, E. Collegial Phenomenon : The Social Mechanisms of Cooperation Among Peers in a Corporate Law Partnership Introduction. in Collegial Phenomenon : The Social Mechanisms of Cooperation Among Peers in a Corporate Law Partnership 346 (Oxford University Press, 2001).

25. Snijders, T. A. B., Pattison, P. E., Robins, G. L. & Handcock, M. S. New Specifications for Exponential Random Graph Models. Sociological Methodology 36, 99–153 (2006).

26. Schlichtkrull, M. et al. Modeling Relational Data with Graph Convolutional Networks. arXiv:1703.06103 [cs, stat] (2017).

27. Paul, S. & O’Malley, A. J. Hierarchical longitudinal models of relationships in social networks. J R Stat Soc Ser C Appl Stat 62, 705–722 (2013).

28. Wasserman, S. S. A Stochastic Model for Directed Graphs with Transition Rates Determined by Reciprocity. Sociological Methodology 11, 392–412 (1980).

29. Veličković, P. et al. Deep Graph Infomax. arXiv:1809.10341 [cs, math, stat] (2018).

30. O’Malley, A. J. & Marsden, P. V. The Analysis of Social Networks. Health Serv Outcomes Res Methodol 8, 222–269 (2008).

31. Krizhevsky, A., Sutskever, I. & Hinton, G. E. ImageNet Classification with Deep Convolutional Neural Networks. in Advances in Neural Information Processing Systems 25 (eds. Pereira, F., Burges, C. J. C., Bottou, L. & Weinberger, K. Q.) 1097–1105 (Curran Associates, Inc., 2012).

32. Kipf, T. N. & Welling, M. Semi-Supervised Classification with Graph Convolutional Networks. arXiv:1609.02907 [cs, stat] (2017).

33. Hamilton, W. L., Ying, R. & Leskovec, J. Representation Learning on Graphs: Methods and Applications. arXiv:1709.05584 [cs] (2018).

34. Smith, A. L., Asta, D. M. & Calder, C. A. The Geometry of Continuous Latent Space Models for Network Data. Statist. Sci. 34, 428–453 (2019).

35. Kingma, D. P. & Welling, M. Auto-Encoding Variational Bayes. arXiv:1312.6114 [cs, stat] (2014).

36. Fard, M. M., Thonet, T. & Gaussier, E. Deep $k$-Means: Jointly clustering with $k$-Means and learning representations. arXiv:1806.10069 [cs, stat] (2018).

37. Karim, M. R. et al. Deep learning-based clustering approaches for bioinformatics. Brief Bioinform doi:10.1093/bib/bbz170.

38. Bianchi, F. M., Grattarola, D. & Alippi, C. Spectral Clustering with Graph Neural Networks for Graph Pooling. arXiv:1907.00481 [cs, stat] (2020).

39. Pope, P. E., Kolouri, S., Rostami, M., Martin, C. E. & Hoffmann, H. Explainability Methods for Graph Convolutional Neural Networks. in 2019 IEEE/CVF Conference on Computer Vision and Pattern Recognition (CVPR) 10764–10773 (2019). doi:10.1109/CVPR.2019.01103.

40. Veličković, P. et al. Graph Attention Networks. arXiv:1710.10903 [cs, stat] (2018).

41. Sundararajan, M., Taly, A. & Yan, Q. Axiomatic Attribution for Deep Networks. arXiv:1703.01365 [cs] (2017).

42. Ying, R., Bourgeois, D., You, J., Zitnik, M. & Leskovec, J. GNNExplainer: Generating Explanations for Graph Neural Networks. (2019).

43. Kipf, T. N. & Welling, M. Variational Graph Auto-Encoders. arXiv:1611.07308 [cs, stat] (2016).

44. O’Malley, A. J., Zou, K. H., Fielding, J. R. & Tempany, C. M. C. Bayesian Regression Methodology for Estimating a Receiver Operating Characteristic Curve with Two Radiologic Applications: Prostate Biopsy and Spiral CT of Ureteral Stones. Academic Radiology 8, 713–725 (2001).

45. Ruopp, M. D., Perkins, N. J., Whitcomb, B. W. & Schisterman, E. F. Youden Index and Optimal Cut-Point Estimated from Observations Affected by a Lower Limit of Detection. Biom J 50, 419–430 (2008).

46. Liu, Y., Li, Z., Xiong, H., Gao, X. & Wu, J. Understanding of Internal Clustering Validation Measures. in 2010 IEEE International Conference on Data Mining 911–916 (2010). doi:10.1109/ICDM.2010.35.

47. Emmons, S. & Mucha, P. J. A Map Equation with Metadata: Varying the Role of Attributes in Community Detection. Phys. Rev. E 100, 022301 (2019).

48. Learning on Graphs: Supervised and Unsupervised Methods.

49. Lee, J., Lee, I. & Kang, J. Self-Attention Graph Pooling. (2019).

50. Rosenberg, A. & Hirschberg, J. V-Measure: A Conditional Entropy-Based External Cluster Evaluation Measure. in Proceedings of the 2007 Joint Conference on Empirical Methods in Natural Language Processing and Computational Natural Language Learning (EMNLP-CoNLL) 410–420 (Association for Computational Linguistics, 2007).

51. Leenders, R. Th. A. J. Modeling social influence through network autocorrelation: constructing the weight matrix. Social Networks 24, 21–47 (2002).

52. Pareja, A. et al. EvolveGCN: Evolving Graph Convolutional Networks for Dynamic Graphs. arXiv:1902.10191 [cs, stat] (2019).

53. Bodnar, C., Cangea, C. & Liò, P. Deep Graph Mapper: Seeing Graphs through the Neural Lens. arXiv:2002.03864 [cs, stat] (2020).

